# NLRP3-caspase 1 and caspase 11-gasdermin D pathways jointly promote IL1β-mediated neuroinflammation and behavioral deficits in a rat model of psychosocial stress

**DOI:** 10.1101/2025.02.02.636090

**Authors:** Soni Tiwari, Zaidan Mohammed, Ekta Yadav, Santi Ranjan Atta, Shweta Kaushik, Aryan Tiwari, Amla Chopra, Itender Singh, Simantini Ghosh

## Abstract

Despite serious health and economic burden, borne more prominently by lower and lower middle-income countries, pharmacotherapy for common psychiatric disorders rely on drugs altering neurotransmission, with partial efficacies. The persistence of a chronic yet sub-threshold inflammation across the periphery and the brain is well documented in the patients and animal models of these disorders and can be investigated for augmenting pharmacotherapy. IL1β, a pleiotropic cytokine and a key regulator of neuroinflammation has been of particular interest in this regard. Previous studies on rodent models of anxiogenic stress show activation of a large multiprotein complex - the NLRP3 inflammasome, which increases IL1β production facilitated by caspase 1, through what is considered a canonical inflammasome activation pathway. However, there exists a second non canonical inflammasome activation pathway whose impact on IL1β release, inflammation and behavioral consequences remain unclear. Using rat models of physical and psychosocial stress, we observed stress-induced sex-specific upregulation of activated caspase 11 and gasdermin D N-terminal fragments, the mediators of the non-canonical inflammasome pathway that facilitates IL1β release through pore formations in the plasma membrane. This is the first report of non-canonical inflammasome pathway being activated in the brain and peripheral immune cells in response to psychosocial stress. Inhibition of caspase 11 with wedelolactone, or Gasdermin D cleavage with Disulfiram reduced stress-induced elevation of IL1β levels, anxiety and fear acquisition, facilitated fear extinction and recall, and improved working memory. Combination treatments targeting both canonical (ibrutinib for pBTK/NLRP3 or MCC950 for NLRP3 inhibition) and non-canonical (wedelolactone for caspase 11 or disulfiram for gasdermin D) pathways proved more efficacious in reducing stress-mediated neuroinflammation, dendritic spine elimination in the CA3 region of the hippocampus and behavioral dysfunction. Furthermore, psychosocial stress drove peripheral inflammation, in peripheral blood mononuclear cells, which was mitigated by the combination treatment. Taken together, this study reveals a novel mechanism underlying psychosocial stress involving both canonical and non-canonical inflammasome signaling to facilitate IL1β induction and behavioral changes. Our finding further suggests that combined targeting of the NLRP3 inflammasome and gasdermin D could contribute to the development of future transdiagnostic therapeutic targets for stress, anxiety, and depression.

## Introduction

The Global Burden of Disease (GBD) study conducted by the World Health Organization (WHO) assessing the burden of psychological disorders worldwide reported 15% of life years lost to these disorders [1]. However, when the definition is expanded, the epidemiological and economic burden of psychological disorders between 2000 and 2019 are even larger and is estimated at 418 million disability-adjusted life years (DALYs) and an economic burden of about USD 5 trillion [2]. Common psychiatric disorders related to anxiety, depression, and traumatic stress constitute a larger proportion of this burden, which is disproportionately borne by lower- and middle-income countries [1,3,4].

Additionally, anxiety-related disorders in general have a much earlier onset compared to severe psychological disorders [5]. The treatment gap is also significantly large in countries of the global south, including India [6,7]. Nevertheless, common psychiatric disorders remain under-researched [8]. The primary pharmacological therapy for individuals with these disorders is limited to selective serotonin reuptake inhibitors (SSRIs) and drugs that alter neurotransmitter levels in the brain, yet only have partial efficacy [9–15]. The situation has been worsened globally by the pandemic, but predictably the lower-middle-income countries have once again borne the brunt of the exacerbation [16]. Considering this, there is a need for a scalable and feasible transdiagnostic approach towards pharmacotherapy across common psychiatric disorders. SSRIs are the usual choice for most stress and anxiety-related disorders [17,18]. Meta-analyses indicate that SSRIs are linked with small to moderate effect sizes in anxiety disorders, post-traumatic stress disorder (PTSD), and major depressive disorder (MDD) [19–21], and are also linked with side effects despite their popularity [22–24]. Given the scale of the problem posed by common psychiatric disorders and the substantial gap, there is a considerable potential to investigate therapeutic strategies that extend beyond neuronal and neurotransmitter-related correlates. Of particular interest here is the extensive evidence showing that stress, anxiety, and depression are characterized by a chronic yet low-grade inflammation, as demonstrated in human and animal studies. Immune dysregulation and elevated levels of proinflammatory cytokines such as IL1β, TNFα, and IL6 are closely correlated with PTSD, anxiety, and depression-like disorders [25–28]. The induction of these cytokines is also strongly associated with symptom severity in both anxiety and depression [29].

Overarchingly, we are interested in investigating therapeutic strategies that modulate these cytokines, specifically IL1β, a major proinflammatory cytokine in the brain, and observing their impact in experimental *in vivo* models. IL1β is known as a key mediator of neuroinflammation and pathophysiology in neurodegeneration, traumatic brain injury, and stroke [30–34]. IL1β induces the expression of multiple other genes across various cell types, including monocytes and macrophages, and promotes the expression of its own genes, creating a positive feedback loop [35–37].

It is in this context that we focus on the NOD-, LRR-, and pyrin domain-containing protein 3 (NLRP3) inflammasome, a multiprotein complex that is an evolutionarily important sensor of pathogen-associated molecular patterns (PAMPs) and damage-associated molecular patterns (DAMPs) of the innate immune system. It also serves as an upstream regulator of IL1β production. NLRP3 inflammasome is characterized by an oligomeric structure, maintained in an inactive state via interactions among the C-terminal LRR domains. In response to specific stimuli, NLRP3 assembles with apoptosis-associated speck-like protein (ASC) and forms a supramolecular complex known as the inflammasome, which activates caspase 1, resulting in the secretion of IL1β [38]. The assembly of the NLRP3 inflammasome leads to caspase 1-dependent induction of the pro-inflammatory cytokines IL1β and IL18, along with gasdermin D (GSDMD)-mediated pyroptotic cell death [39]. IL1β is produced by the cysteine protease caspase 1 following the cleavage of its precursor protein pro-IL1β, marking the final stage of the NLRP3 inflammasome activation [40]. Aberrant activation of the NLRP3 inflammasome is associated with the pathogenesis of multiple inflammatory diseases, including diabetes, lifestyle conditions such as smoking, cancer, and Alzheimer’s disease [38].

Recently, the NLRP3 inflammasome has also been attracting attention in psychological disorder research [41–43]. The NLRP3 inflammasome is elevated in the brain and blood of depressive patients [43,44]. It is a key mediator of pro-inflammatory activity and the release of IL1β in anxiety and depression [45–47]. We have previously demonstrated that Bruton’s tyrosine kinase (BTK) acts as an upstream regulator of NLRP3 activation in mouse models of physical restraint stress. Stress-induced activation of BTK upregulates IL1β and other neuroinflammatory mediators, leading to exacerbated anxious behavior. We also demonstrated that BTK inhibition with ibrutinib led to amelioration of this heightened anxious phenotype with a concomitant reduction in NLRP3, IL1β and neuroinflammation [48]. In this article we extend our studies to investigate a second, non-canonical inflammasome pathway that contributes to the release of IL1β from cells.

Canonical NLRP3 inflammasome activation leads to the activation of caspase 1 that cleaves pro-IL1β into IL1β. However, caspase 1 can also cleave GSDMD, a 480 amino acid long protein, into an N-terminal fragment (GSDMD-N). These fragments assemble to form small transmembrane pores, which facilitate the release of more IL1β into the interstitial space. The increase in the number of these pores can disrupt the ion and water balance, exacerbating inflammation through a process called pyroptosis [49]. Apart from the canonical activation of caspase 1, there is also a non-canonical inflammasome pathway through caspase 11. Mechanistically, caspase 1 and caspase 11 specifically cleave the linker between amino-terminal and carboxy-terminal domains of GSDMD, which is required and sufficient for the pore formation in the membrane [50,51]. The cleaved N-terminus can undergo auto-oligomerization on membranes upon interaction with phosphoinositides or other acidic lipids, leading to the creation of small circular membrane pores of around ∼21 nm in diameter [52,53]. This pore formation helps with the release of 17 kDa IL1β from the cells [54]. GSDMD-mediated pyroptosis plays a key role in cell death and host survival, and its dysregulation may contribute to cancer progression [55–57]. Lipopolysaccharide signalling leads to profound activation of caspase 11 and its human counterparts caspase 4/5, which leads to cleavage of GSDMD and contributes to the downstream sequelae as described above [50,58]. Initially limited to bacterial infections, caspase 11-mediated pyroptosis has been associated with various disease pathologies, such as lung injury and asthma, skin psoriasis, peptic ulcers in gastric tissue, and graft versus host disease [59–62]. Recent studies in chronic stress-induced depression models have explored the role of canonical caspase 1-mediated GSDMD pathway in neuroinflammation [63–65]. However, the role of non-canonical activation by caspase 11 under psychosocial stress has not been explored yet.

In this study, utilizing rat models of stress, we investigate both canonical and non-canonical inflammasome pathways leading to IL1β release and associated anxiety-like behavior, fear conditioning and working memory deficits. Building upon our earlier work on mouse models with stress-induced activation of canonical NLRP3 inflammasomes, we further extend our investigation into non-canonical activation of caspase 11 and gasdermin D in rats, which arguably have a wider repertoire of complex social behaviors that mice can exhibit and are a more advanced model of studying anxiety and cognitive behaviors. Additionally, concomitant to a physical stress model, we also show that inducing psychological stress by exposing rats to a repeated social defeat paradigm leads to significant neuroinflammation, neuronal deficits, and prominent behavioral dysfunction across various domains. This allows us to get a closer approximation of human stress experience, as the social defeat paradigm is regarded as one of the most indicative animal models for investigating social stress [66]. We then apply known inhibitors of the mediators of the canonical NLRP3 inflammasome and non-canonical caspase 11-GSDMD activation, both individually and in combination, to stressed rodents. We demonstrate that a combination treatment approach offers greater efficacy to improve neural correlates, mitigate neuroinflammation, and alleviate behavioral deficits in the psychosocial stress model. Among the inhibitors we used, ibrutinib and disulfiram are approved by the FDA for treatment of cancers and alcohol use disorder, respectively, thereby keeping our work relevant from a translational perspective [67–71]. The primary contribution of this study is the discovery of previously unidentified stress-induced activation of non-canonical inflammasome mediators that drives neuroinflammation, dendritic spine elimination, and several dimensions of behavior deficits. Another key finding of this study from the pharmacological inhibitor experiments is that the simultaneous inhibition of both canonical and non-canonical inflammasomes in stressed animals is more beneficial than individually inhibiting these pathways.

## Methods

### Experimental animals

Male and female Sprague Dawley (SD) rats were obtained from the National Institute of Biologicals, Noida, India, and the All India Institute of Medical Sciences, New Delhi, India. The rats were group housed in cohorts of 3 per cage with *ad libitum* access to rodent chow and filtered water. The rats were kept in a vivarium under standard conditions (21°C ± 2°C temperature, 55% ± 10% humidity, a 12 h:12 h light-dark cycle starting at 7 am, 20–22 lux light inside the cage, and corn cob bedding).

Rats were acclimatized for at least two months before the start of the experiments. All animal experiments were conducted according to the guidelines and protocols approved by the Animal Ethics Committees of Delhi University, India, and Maharshi Dayanand University, Rohtak, India. Physiological parameters such as temperature, heart rate, and body weight were monitored during the experiments. Experiments were initiated at 5 months of age.

### Pharmacological inhibition of inflammasomes

Five-month-old SD rats were randomly assigned to control and different drug treatment groups. Rats received canonical and non-canonical inflammasome inhibitors or vehicles, starting at 5 h before the induction of stress, and again given every 24 h for a total of 7 days. To pharmacologically inhibit caspase 11, wedelolactone dissolved in dimethyl sulfoxide (DMSO): polyethylene glycol 400 (PEG): water at 1:4:5 ratios was administered by oral gavage (30 mg/Kg/day; Sigma-Aldrich, St. Louis, MO), as reported earlier [72]. Control rats were similarly dosed intragastrically (i.g.) with vehicle (DMSO: PEG). To inhibit gasdermin D, rats were administered intraperitoneally (i.p.) with disulfiram (50 mg/Kg/day; Sigma-Aldrich) formulated in sesame oil (12.5 mg/ml), as reported earlier [68]. Control animals were similarly dosed with vehicle (sesame oil, i.p.). To validate the observations from disulfiram studies, another gasdermin D inhibitor, necrosulfonamide (Sigma-Aldrich) dissolved in DMSO, was also used in a separate set of animals (20 mg/Kg/day, i.p.), as reported earlier [73].

Control rats were similarly injected with vehicle (0.05% DMSO in saline). To pharmacologically inhibit NLRP3 inflammasome, a specific inhibitor MCC950 (50 mg/Kg/day, i.p.; Sigma-Aldrich) dissolved in DMSO was used, as we reported earlier [48]. Control rats were injected with 0.05% DMSO in saline (i.p). To inhibit the upstream pathway of the NLRP3 inflammasome, a Bruton’s tyrosine kinase (BTK) inhibitor ibrutinib (3 mg/Kg/day; i.p.; Sigma-Aldrich), dissolved in DMSO, was used, as we reported earlier [48]. Control rats were injected in parallel with the vehicle (0.05% DMSO in saline) at the indicated times. To determine the effect of simultaneously inhibiting canonical and non-canonical inflammasomes, we administered MCC950 (50 mg/Kg/day; i.p.), a canonical NLRP3 inflammasome inhibitor, and wedelolactone (30 mg/Kg/day, i.g.), a non-canonical inflammasome inhibitor (caspase 11 inhibitor) [74]. To determine the potential therapeutic efficacy of simultaneous inhibition of canonical and non-canonical inflammasomes, we also used FDA-approved inhibitors, ibrutinib (3 mg/Kg/day; i.p.) and disulfiram (50 mg/Kg/day; i.p.).

### Stress paradigms

We used two different types of paradigms to mimic stress and induce accompanying behavioral changes in 5-month-old SD rats. Rats were randomly assigned to control and stress conditions. Restraint stress followed by underwater submersion was used for the physical stress paradigm. Repeated social defeat was used to induce psychosocial stress. Three weeks prior to behavior experiments, rats randomly assigned to either control or stress condition were handled daily at the testing arena to reduce the external anxiety. All the behavior procedures were conducted during the dark phase (7 pm and 11:30 pm) of the cycle to avoid the potential influence of daytime noise and disturbances. The stressed animals were housed individually for the rest of the experiment to eliminate the potential influence of social housing on recovery from stress induced behavioral alterations.

### Physical stress

Rats were subjected to physical stress-induced anxious behavior based on a method adapted from our earlier study [48]. In brief, rats were transported from the colony room to a procedure room where the physical stress paradigm was administered. On the first day, rats were restrained by immobilizing them in perforated polycarbonate chambers (measuring 21 cm long x 9 cm wide x 7 cm high) for 3 hours. The restraining chamber was placed in the inclined plane of a 75-degree angle with the head positioned downward during the entire 3 hours of immobilization. Rats were then moved to solitary housing in a separate cage with food and water. The next day, rats were subjected to underwater trauma by first allowing them to swim for 45 s in a container filled with ice-cold water and then suddenly pushing them down to gently submerge them completely underwater for 30 s. Rats were again moved to solitary housing with food and water and housed there for the remaining duration of the experimental procedures. To eliminate any odour cues, the restraining chambers and water containers for underwater trauma were cleaned with a 70% ethanol solution and completely dried before putting in the next rat.

### Psychosocial stress

Five-month-old dominant and aggressive male SD rats were identified and paired with females, with one male and one female in each cage. After a month, these dominant male rats were conditioned for heightened aggression, once per week for 45 days, to attack the intruder by removing the resident female for an hour and placing a new 4-month-old male or female rat as an intruder for 15 minutes. After analysing the conditioning sessions, the most aggressive and dominant male rats were selected for the experiments. To induce psychosocial stress, 5-month-old male and female SD rats randomly assigned to the psychosocial stress condition were used as intruders. One hour before the induction of psychosocial stress, the resident cage mate female rat was removed from the cage containing the aggressor male. A new intruder test rat (male) was placed into the home cage of the aggressor male for 15 minutes. The aggressor rat displayed a full repertoire of aggressive behavior, such as sideway threats, aggressive pursuits, and bites on the intruder, which showed defensive and submissive postures, as described earlier [75]. To prevent further physical injury while allowing the sensory interaction to stimulate the psychosocial threat of attack, the intruder was taken out and put into a small wire mesh cage (25 cm x 10 cm x 10 cm) and then placed back into the home cage of the aggressor, which continued trying to attack the test rat through the mesh. After having spent a total of 3 hours, the defeated intruder rat was taken out and moved to solitary housing, and the aggressive rat was reunited with its female companion in the home cage. This intruder test rat was again subjected to repeated social defeat by new aggressor rats for the next two days. To avoid habituation, different aggressors were used every day to instil psychosocial stress into the intruder test rats. For female rats, the gender of the aggressor was matched with the intruder, and an identical protocol as described above was followed. During the entire course of repeated social defeat, the physical health status of the intruding test rats was carefully monitored, and significantly wounded rats were removed from the study. Only 7% of the rats met the removal criterion. The control animals were simply handled in the colony room and kept with their companion resident rats.

### Analysis of anxiety, fear memory, social interaction and working memory

Seven days after the induction of physical and psychosocial stress, the test rats were assessed for anxiety-like behavior using the open field test, light-dark test, elevated plus maze test, and social interaction tests. Working memory was analyzed by Y-maze and novel object recognition tests. We also tested fear acquisition, extinction, and recall using a fear conditioning paradigm. All trial sessions were video recorded and analyzed by a researcher blinded to the group status of the rats. Anxiety-like behavior and memory function assessment were performed between 7 pm and 11:30 pm during the dark phase of the light cycle when rats are most active, as they are nocturnal and to avoid disturbance from daytime activities. In between each test animal and trials, all behavioral tools and apparatus were thoroughly cleansed and deodorized with 70% ethanol and dried to eliminate olfactory cues from other experimental animals. Four days before the onset of behavior assessment, the animals were shifted to the behavior testing room for 2 hours and remained undisturbed in their home cage to enable them to acclimatize to the new environment. Behavior tests were performed in the following sequence:

### Open field test (OFT)

Analysis of anxious behavior was performed using an open-air (without ceiling) box-shaped apparatus made of black polycarbonate sheets (length 80 cm x width 80 cm x height 60 cm). The open field test was conducted to analyze anxious behavior based on a method adapted from our earlier study [48]. In brief, the test rat was placed in the corner of the empty open field test apparatus and allowed to freely explore it for 5 minutes. The time spent in the central area (40 cm x 40 cm) was measured to assess anxiogenic behavior. The more time spent in 20 cm wide periphery (thigmotaxis) indicated the severity of anxiety. The higher thigmotaxis correlates with the higher level of anxiety-like behavior in rodents, and stressed animals tend to avoid the central area of the open field arena [76].

### Elevated plus maze test (EPM)

The elevated plus maze raised to a height of 100 cm was used to measure anxiety-like behavior related to unsafe height. The plus-shaped platform made of polycarbonate sheets consisted of four arms (60 cm long, 15 cm wide), with two opposite sets of arms enclosed by 30 cm high walls and two other arms remaining open. The opposite arms with enclosed walls represented the secure areas, while the other set of open arms without walls represented the unsafe areas [48]. The arms were brightly illuminated with 100 lux light. To test anxiety-like behavior in rats, each rat was placed in the central square area of the elevated plus maze and allowed to explore freely in all arms for 5 min. The percentage of time spent on the open arm (unsafe area) was recorded to assess the anxious behavior [77]. The rat was considered inside the open arm when all four limbs entered the open arm. Lesser time spent in the open arm was taken as an indicator of a higher degree of anxiety in rats.

### Light-dark test (LDT)

The light-dark test was used to examine the innate tendency of rats to avoid brightly lit areas and move to the safety of dark areas, especially during stress [78]. The box-shaped apparatus consists of two compartments (25 cm length x 25 cm wide x 45 cm height), one covered with black opaque polycarbonate sheets to serve as a dark area and the other with clear polycarbonate sheets to act as a light area. The apparatus was divided into two equal-sized compartments by a barrier that contained an open door (10 cm x 10 cm) to allow the test rat to freely escape from the brighter area to the darker area for safety, especially when stressed. Rats were allowed to explore the light-dark apparatus for 10 min, and relative time in the dark area to the light area was counted [48]. A higher dark/light ratio reflected higher anxiety.

### Three-chamber social interaction test

Social impairment induced by stress negatively impacts both males and females [79]. The psychosocial stress-induced social impairment was analyzed by using the sociability and social preference tests. The three-chamber test apparatus was made up of a polycarbonate box (length 120 cm x width 40 cm x height 40 cm), consisting of three chambers. The middle chamber contained clear polycarbonate sliding windows toward both side chambers to enable sniffing by rats. A removable top lid was used to place the animals in the respective chambers. First, the test rat was habituated by letting it explore the middle chamber with fully opened side windows for 10 min and then transferred back to the home cage. A new rat, referred to as rat 1 of the same age and gender as the test rat, was placed into one of the side chambers with a partially closed transparent polycarbonate window for 10 minutes. Thereafter, for the sociability session, the test rat was placed into the middle chamber and allowed to freely explore for 10 min. Time spent by the test rat attempting to sniff and interact with rat 1 versus the empty chamber on the other side was recorded. The sociability preference index was determined by dividing the time spent as described above by the test rat with rat 1 by the time spent with the empty chamber. At the end of this session, the test rat was removed and placed in the home cage for 10 min. Meanwhile, a novel rat referred to as rat 2 of the same age and gender was placed into the empty side of the chamber. To determine the social preference for the novel rat, the test rat was again placed back into the central chamber for another 10 min. The social preference time was calculated by dividing the time spent by the test rat with the novel rat (rat 2) by the time spent with the familiar rat (rat 1). Both rats 1 and 2 were of the same gender as the test rat. The placement of familiar rats and novel rats was systemically altered between trials to eliminate any chamber bias.

### Novel object recognition test (NOR)

To investigate working memory in anxious rats, a novel object recognition test was performed as previously described [80], with minor modifications. The test rat was first habituated for 30 min to the test apparatus, a black polycarbonate box (length 80 cm x width 80 cm x height 60 cm). Thereafter, the test rat was removed and put back in its home cage for 20 min. Two identical plastic toys (same size, shape, colour and texture) were placed approximately 10 cm away from the diagonally opposite corners of the test chamber. In the first familiarization phase, the test rat was allowed to freely explore the two identical objects for 10 min. At the end of this session, the test rat was again removed and returned to its home cage. Meanwhile, one of the familiar test objects was replaced with a non-identical novel object (different size, shape, colour and texture). In the final phase of NOR, the test rat was allowed to explore for 5 min with two of these objects, one familiar and another novel object. The exploration time spent by the test rat with familiar and novel objects was measured. The percentage of exploratory preference for novel object was determined to evaluate the anxiety-induced working memory deficits.

### Y-maze test

To assess the spatial working memory, we used the Y-maze test to determine the number of entries to the novel unexplored arm, as previously described [81]. The black polycarbonate Y-maze apparatus consisted of three identical arms projected in a Y-shape (length 60 cm x width 15 cm x height 30 cm). Visual-spatial cues were provided around the Y-maze by placing differently coloured papers and objects on the top side of the walls and roof of the behavioral testing room to help differentiate the relative position of the three arms. In the first phase of the test, the entry to one of the three arms was blocked with a black polycarbonate removable door. In this phase, the test rat was placed at the start of one of the open arms and allowed to explore for 15 min, which is considered an acquisition trial. Rats were allowed to familiarize themselves with these two open arms to acquire the spatial cues and differentially identify the arms. During the entire period of the acquisition trial, the entry to the novel arm remained closed. At the end of acquisition trial, the test rat was removed and placed in home cage for 30 min. During the test trial, the entry to the novel arm was freely allowed by completely removing the black polycarbonate door. Rats were allowed to explore all three arms of the Y-maze for 15 min in this phase, referred to as the recall trial. The number of entries to the novel arm by this test rat was counted and compared to the total entries in all three arms. A normal unstressed rat preferably explores the novel arm more as compared to the two familiarized arms, indicating its ability to recall previously visited arms. Anxiety is associated with impairment of spatial memory, and anxious rats show reduced preference to the novel arm.

### Fear conditioning tests

The Pavlovian fear conditioning tests were performed to assess fear memory, as previously described [82], with minor modifications. In brief, we measured fear acquisition, extinction, and recall over three consecutive days using polycarbonate chambers (length 45 cm x width 24 cm x height 40 cm). The apparatus was equipped with a speaker that emitted a 3000 Hz tone amplified to 86 dB. The floor of the chamber was made of a stainless-steel grid to deliver a foot shock (500 ms, 0.5 mA). A day before the fear acquisition test, the rats were acclimatized to the fear conditioning apparatus by placing them in the chamber for 30 minutes without any sound or foot shock. On the first day of the fear acquisition trial, the test rat was allowed to freely move in the test chamber for 5 min of habituation. Thereafter, the test rat was subjected to five fear conditioning trials for fear acquisition. Each conditioning trial for fear acquisition consisted of 20 sec tones from the speaker (conditioned stimulus) and a foot shock (unconditioned stimulus) in the last 2 sec of the culmination of sound. Freezing time, defined by the absence of any visible movement except for respiration-related movement, was measured during the 20 sec of conditioned sound. Each trial was separated by 210 sec intervals. The test rat was moved back to the home cage after the fear acquisition trial. On the second day, the test rat was again placed in the fear conditioning chamber and allowed to habituate for 5 min, followed by 10 fear extinction trials using only the conditioned stimulus, i.e., 20 sec of the tone without any foot shock. To determine fear extinction, freezing time was similarly recorded during these 20 secs of conditioning stimulus. The inter-trial period was 210 sec, during which no stimulus was provided. The test rat was then moved back to its home cage. On the third day, after 5 min of habituation, the test rat was subjected to 10 trials of fear recall by exposing it to only a conditioned stimulus of 20 sec sound, without any foot shock. The inter-trial interval in the fear recall trial was also 210 sec. Freezing behavior was similarly recorded over these 10 fear-cued recall trials. Percent freezing time (sec spent freezing/20 sec conditioned stimulus) during fear acquisition, extinction, and recall was calculated for stressed and control rats.

### Tissue collection

Serial blood samples to analyze corticosterone levels at various time points before and after the induction of stress were collected from the jugular vein, as previously described [83]. In brief, a catheter was inserted into the right jugular vein of the rats and connected to polyethylene tubing (PE-50) with heparinized saline (0.9% NaCl, 25 IU/ml). Blood samples of 150 µl were withdrawn using a syringe and centrifuged at 2000 g for 30 min at 4°C to collect plasma. Samples were stored in a −80°C freezer for later use. To collect other samples at eleven days after the induction of stress, rats were anesthetized with ketamine (80 mg/Kg) and xylazine (10 mg/Kg). First, 150 µl of cerebrospinal fluid (CSF) was carefully collected from the cisterna magna, while avoiding contamination from blood, as described earlier [84]. Thereafter, about 3 ml of blood was collected directly via cardiac puncture and centrifuged to collect the plasma. The rats were transcardially perfused with 200 ml of ice-cold phosphate-buffered saline (PBS) to maximally clear blood from the brain vasculature. The brains were quickly removed, and various regions of the brain, like the hippocampus, amygdala, and prefrontal cortex, were carefully yet rapidly dissected under stereomicroscope, while on ice and immediately stored in a −80°C freezer for later use.

### Golgi-Cox staining for dendritic spines

Golgi-Cox staining was performed to analyze apical dendritic spine density and the number of mushroom-shaped spines in the cornu ammonis 3 (CA3) region of the hippocampus in both stressed and control rats. The Golgi-Cox method consists of various steps, which include specimen impregnation with Golgi-Cox solution, cryoprotection, sectioning, colour development, and mounting, as described earlier [85,86]. In brief, rat brains were isolated and impregnated in the dark with a Golgi-Cox solution containing potassium dichromate, mercuric chloride, and potassium chromate for 40 days at room temperature. To preserve tissue morphology, brains were transferred into a cryoprotectant solution containing 20% sucrose and 15% glycerol in water for 3 days. Coronal rat brain sections of thickness 100 μm were cut using a vibratome (VT1200S, Leica, Nussloch, Germany). Thereafter, sections were mounted on chrome alum-gelatinized slides. During the colour development step, slides were washed with double-distilled water and then incubated in a 75% ammonia solution for 10 min in the dark. Sections were washed with water and fixed by treatment with a 1% sodium thiosulphate solution for 10 min. The brain sections were dehydrated with ethyl alcohol series (70%, 90%, and 100%) and cleared in xylene. Subsequently, sections were mounted using Permount mounting medium, and slides were kept in the dark until microscopic examination. To visualize dendritic spines, brightfield images with a frame size of 1024×1024 pixels were captured using a 100x objective with a 1.3 numerical aperture and a BX63 Olympus microscope (Olympus Corporation, Tokyo, Japan). Five different sections from each rat were used to determine the mean spine density and mean number of mushroom spines per 10 µm of apical dendrite in the CA3 region of the hippocampus. The dendrites directly emerging from the primary shaft from the soma were classified as primary apical dendrites, as described earlier [87]. Dendritic spines from the CA3 region were categorized and counted with the open-source software ImageJ (National Institute of Mental Health, Bethesda, MD). Mushroom spines were classified by their shape of mushroom, with a head diameter greater than 0.5 µm and a head-to-neck ratio greater than 1:1. The number of spines was counted by an experimenter blinded to the identity of the samples.

### Isolation and culture of peripheral blood mononuclear cells

Rat peripheral blood mononuclear cells (PBMC) were isolated from the 3 ml of heparinized blood collected through cardiac puncture. The separation and isolation of PBMCs from rats were performed by density gradient centrifugation using OptiPrep solution (Sigma-Aldrich), as we described earlier [48]. In brief, OptiPrep density gradient solution of iodixanol in water (60% w/v) was layered with diluted blood in buffered saline (0.85% NaCl, 10 mM HEPES, pH 7.4) and centrifuged at 400 g for 30 minutes at 20°C. PBMCs were carefully collected from a sharp band at the interface and washed twice with tricine-buffered saline. The cells were pelleted by centrifugation (200 g, 15 minutes at 20°C).

The purity of PBMC, determined morphologically by May-Grünwald Giemsa (MGG) staining, was nearly 99%. To culture, PBMCs were resuspended in RPMI 1640 medium (Sigma-Aldrich) supplemented with 10% fetal calf serum (Hyclone, Logan, UT). PBMC were cultured to the concentration of 2.5×10^6^ cells/ml, and a set of these cells was primed with lipopolysaccharides (LPS) from *Escherichia coli* serotype 0111: B4 (100 ng/ml) and nigericin (10 µM). The control and primed cells were treated with various inhibitors of inflammasomes, such as wedelolactone (40 µM), MCC950 (5 µM), ibrutinib (1 µM), disulfiram (100 nM), or 0.2% DMSO as vehicle control. The cell culture supernatant and cells were collected and kept in a −80°C freezer for later use.

### Cell viability assay

The viability of PBMCs was determined using the WST-8 assay, as described earlier [88]. Briefly, non-toxic 2-(2-methoxy-4-nitrophenyl)-3-(4-nitrophenyl)-5-(2,4-disulfophenyl)-2H-tetrazolium monosodium salt (WST-8) (Dojindo Molecular Technologies, Gaithersburg, ML) was added to the control and LPS-primed PBMCs treated with inhibitors. PBMCs were incubated in a cell culture CO_2_ incubator for an hour to allow cells to metabolize WST-8 to form a water-insoluble formazan product, which is detected by absorbance at 460 nm. The absorbance of WST formazan at 460 nm is directly proportional to the cell viability.

### Cell permeabilization assay

SYTOX Green was used to measure cell permeabilization, as described previously [89]. SYTOX Green is a low molecular weight (<0.6 kDa) DNA-binding fluorescent dye that infiltrates into cells following plasma membrane pore formation, an indicator of cell pyroptosis. SYTOX Green is not permeable to undamaged cells. The fluorescent intensity of SYTOX Green increases more than 500-fold following binding with DNA, making the onset of pyroptosis/pore formation more sensitively detected based on its uptake. PBMCs were treated with vehicle, LPS, and inhibitors as indicated earlier. To quantify cell permeabilization, cells were treated with 2.5 mM SYTOX Green (Sigma-Aldrich), and cells were kept in a CO2 incubator. To determine cell permeabilization, fluorescence at 528 nM after excitation at 485 nM was recorded using a Biotek Synergy plate reader (Agilent, Santa Clara, CA).

### Immunoblot analysis

The rat brain tissue samples and PBMCs were analyzed for the protein levels of gasdermin D N-terminal fragment, cleaved caspase 11, NLRP3, cleaved caspase 1, and β-actin by immunoblotting, as we described earlier [48,90]. In brief, brain tissue samples and PBMCs were homogenized in RIPA lysis buffer with protease and phosphatase inhibitor cocktails (Thermo Fisher Scientific, Waltham, MA) using a sonicator (Mesonix Sonicator 3000, Cole-Parmer, Vernon Hills, IL). These homogenates were centrifuged at 14,000 g for 25 min at 4°C, and then protein content was measured by BCA protein assay kit (Thermo Fisher Scientific). Equal protein samples were prepared in Laemmli sample buffer and separated by SDS PAGE electrophoresis (Bio-Rad Laboratories, Hercules, CA) on 4-12% Tris-glycine gels (4-12%). The proteins were transferred onto the PVDF membrane (Bio-Rad Laboratories) and blocked with 5% bovine serum albumin (IgG-free endoprotease-free BSA, Jackson ImmunoResearch Laboratories, West Grove, PA) in 50 mM Tris–HCl (pH 7.4), 150 mM NaCl, and 0.1% Tween 20 (Tris-buffered saline with Tween-20, TBST). The PVDF membranes were incubated for overnight at 4°C with the following primary antibodies: anti-cleaved gasdermin D (Asp276, rabbit monoclonal antibody, 1:1000, #10137, Cell Signaling Technology, Danvers, MA); anti-cleaved caspase 11 p20 (mouse monoclonal antibody, 1:1000, #50-256-3377, Fisher Scientific, Waltham, MA); anti-NLRP3 (D4D8T, rabbit monoclonal antibody, 1:1000, #15101, Cell Signaling Technology); anti-cleaved Caspase 1 (p20, Asp296, E2G2I, rabbit monoclonal antibody, 1:1000, #89332, Cell Signaling Technology), β-actin (mouse monoclonal antibody, 1:10000, #sc-47778, Santa Cruz Biotech, Dallas, TX). The membranes were washed five times with TBST and incubated for 1 h at room temperature with horseradish peroxidase-conjugated goat-anti rabbit IgG secondary antibody (1:10,000, #STAR124P, Bio-Rad Laboratories) or horseradish peroxidase-conjugated goat anti-mouse IgG (1:10,000, #sc-2005, Santa Cruz Biotech). Membranes were washed again with TBST, and protein bands were detected using SuperSignal West Pico PLUS Chemiluminescent substrate (Thermo Fisher Scientific).

### Quantitative real-time PCR

The hippocampal samples from experimental rats were used to determine the mRNA levels of C-C motif chemokine ligand (CCL2), CC chemokine receptor 2 (CCR2), and 18S as a housekeeping gene by quantitative reverse transcription polymerase chain reaction (qRT-PCR). The total RNA from the hippocampal samples was extracted by RNeasy kit and treated with DNase (Qiagen, Germantown, MD) following manufacturer’s instructions. The concentration of RNA preparation was assessed at 260 nm. Samples with absorbance ratios of 260/280 and 260/230 greater than 1.7 were used for used for qRT-PCR. RNA samples were reverse transcribed by using SuperScript III reverse transcriptase (200 U) and poli-T primer (0.1 µg, Invitrogen, Carlsbad, CA). The target cDNA was amplified using HotStart-IT SYBR Green qPCR master mix (USB, Cleveland, OH) and the RT-PCR system (Applied Biosystems, Foster City, CA). Following oligonucleotide primers were used *CCL2 F-AGCCAACTCTCACTGAAGC, CCL2 R-GTGAATGAGTAGCAGCAGGT, CCR2 F-CACCGTATGACTATGATGATG, CCR2 R-CAGGAGAGCAGGTCAGAGAT, 18S F-AGTCGCCGTGCCTACCAT* and *18S R-GCCTGCTGCCTTCCTTG*. The relative mRNA levels were calculated using the comparative threshold cycle CT (ΔΔCT) method, as described previously [91].

### Caspase 1 activity assay

Caspase 1 enzymatic activity in rat brain samples and PBMC was assessed using a caspase 1 colorimetric assay kit based on spectrophotometric detection of chromophore p-nitroaniline (p-NA) after cleavage from substrate YVAD-pNA, as per the manufacturer’s instructions (YVAD, Merck Millipore, Burlington, MA). In brief, the brain tissue and PBMC were homogenized in ice-cold lysis buffer and mixed with 2.5 nM dithiothreitol and 50 µl caspase 1 reaction buffer in a 96-well flat-bottom microplate. A substrate solution containing YVAD-pNA was added to each well, followed by incubation at 37°C for 2 h. The enzymatic activity of caspase 1 was measured at 405 nm using a plate reader (Molecular Devices, San Jose, CA).

### Enzyme-linked immunosorbent assays (ELISA)

The rat brain, plasma, CSF, and PBMC samples were used to assess the levels of corticosterone, IL1β, gasdermin D, C-X-C motif chemokine ligand 2 (CXCL2), and intracellular adhesion molecule-1 (ICAM1) using commercially available enzyme-linked immunosorbent assays (ELISA) kits, as per manufacturers’ instructions. In brief, the plasma levels of corticosterone were measured by DetectX corticosterone ELISA kits (#K014-H1, Arbor Assays, Ann Arbor, MI). IL1β levels in rat brain, plasma, and PBMC were quantified by IL1beta Quantikine ELISA kits (#RLB00, R&D Systems Inc, Minneapolis, MN). Gasdermin levels in the rat brain CSF were measured by GSDMD ELISA kits (#orb1784611, Biorbyt, Durham, NC). Rat hippocampal samples were used to assess the levels of CXCL2 using the CXCL2/CINC-2 Quantikine ELISA kit (#RCN300, R&D Systems Inc). ICAM1 levels in rat hippocampus were measured by using the ICAM1/CD54 Quantikine ELISA kit (#RIC100, R&D Systems Inc).

### Statistical analysis

For statistical analysis, the open-source program JAMOVI (The Jamovi project 2022, Version 2.3) was utilized. Most of our data needed different factorial ANOVA designs, which are mentioned in relevant result sections below. Given our prior experience in these studies, we postulated that for most of our tests, interaction terms rather than main effects would be significant. Consequently, we have only reported significant interaction terms along with their partial eta square (ηp2) in order to indicate effect sizes, rather than main effects, wherever applicable. In absence of significant interactions, we have reported main effects with effect size. All significant main effects and interactions were also broken down with post-hoc testing or simple main effects analysis as and when required. For ANOVAs that needed more complex design beyond a 2X2 factorial style, the most complex significant interaction term has been analysed post-hoc for pairwise comparison. For each post hoc comparison, t values have been reported in the text alongside p values instead of Cohen’s d as measures of effect size because Cohen’s d in JAMOVI is not adjusted for multiple comparisons. Animal numbers for each experiment are mentioned in the legends for individual figures. A detailed narrative results section for all statistical analysis is included in the supplementary files. The significance threshold has been set at p < 0.05 for all analyses.

## Results

### Both physical and psychosocial stress models elicited a sexually divergent behavioral and neuroinflammatory phenotype in rats

We established two different paradigms of stress, one physical and another psychosocial, in 5-month-old SD rats. The protocol is schematically illustrated in Figure 1A and narrated in detail in the methods section. Both physical and psychosocial stress elicited a robust upregulation of IL1β in plasma, as measured by an ELISA assay (Figure 1B). Our previous experience with mouse models of stress had shown us that female rodents typically display a more aggressive phenotype following stress, both in behavioral and neuroimmune markers. Our present analysis indicated a similar pattern in the rat models as well. We conducted a 2X2 ANOVA to assess the effects of stress type and sex on plasma IL1β levels (ng/ml) at 10 days after the first initiation of stress (Figure 1B). The analysis revealed a significant main effect for stress type [F(2, 64) = 66.30, p < 0.001, ηp²= 0.674], a significant main effect for sex [F(1, 64) = 21.83, p < 0.001, ηp² = 0.254], and also a significant interaction between stress type and sex [F(2, 64) = 6.31, p = 0.003, ηp² = 0.165]. Post hoc pairwise comparisons with Bonferroni correction were conducted thereafter, which showed significant differences in plasma IL1β levels based on stress type and sex. In the control group, no significant differences were observed between female and male rats [M difference = −0.41, SE = 2.00, t(64) = −0.205, p = 1.000]. However, in the physical stress condition, females exhibited significantly higher IL1β levels than males [M difference = 8.62, SE = 2.00, t(64) = 4.30, p < 0.001], and both females [M difference = −19.60, SE = 2.05, t(64) = −9.58, p < 0.001] and males [M difference = −10.57, SE = 1.96, t(64) = −5.40, p < 0.001] had higher levels of IL1β as compared to the control group. Similarly, in the psychosocial stress condition, females had significantly higher IL1β levels than males [M difference = 7.88, SE = 1.96, t(64) = 4.02, p = 0.002], and both females [M difference = −16.99, SE = 2.00, t(64) = −8.48, p < 0.001] and males [M difference = −8.69, SE = 1.96, t(64) = −4.44, p < 0.001] had higher levels of IL1β, as compared to the control group. No significant differences were observed between the physical stress and psychosocial stress conditions for either females [M difference = 2.61, SE = 2.00, t(64) = 1.31, p = 1.000] or males [M difference = 1.88, SE = 1.96, t(64) = 0.96, p = 1.000].

**Figure 1.**
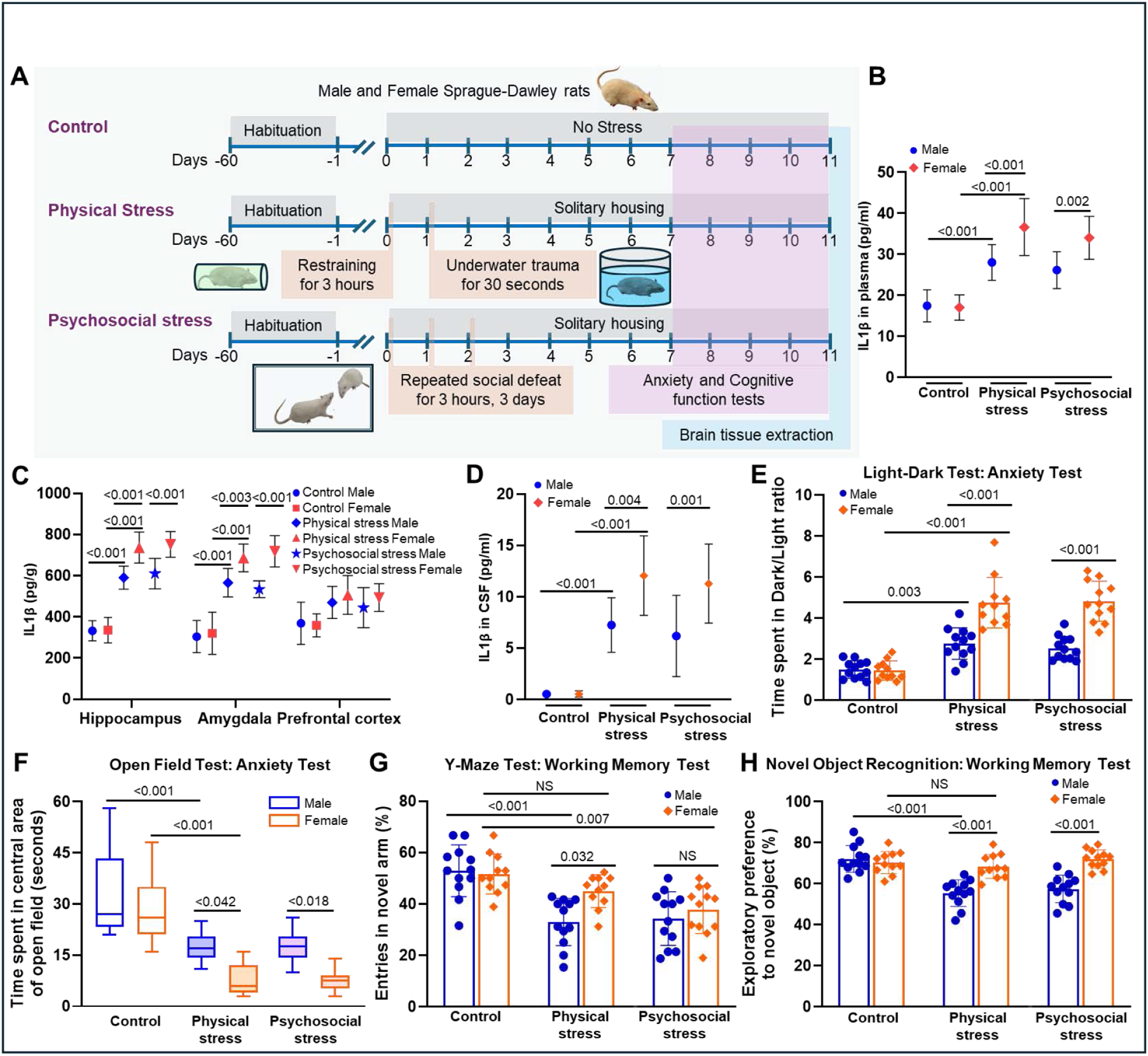
Molecular and behavioral characterization of the rat models of physical and psychosocial stress. **(A)** 5-month-old male and female Sprague-Dawley (SD) rats were used for physical and psychosocial stress models. To elicit physical stress, restraining stress for 3 hours was performed on the first day and then followed by underwater trauma for 30 seconds on the next day. For psychosocial stress, animals were subjected to the social defeat paradigm for 3 consecutive days with different aggressor rats of the same biological sex. Behavioral analysis was performed after one week of stress induction, and all the animals were sacrificed and samples collected after 11 days of stress induction. IL1β levels in plasma (**B**), brain homogenates (**C**), and CSF (**D**) were measured with ELISA. A significantly higher level of IL1β was observed in plasma, CSF, hippocampus, and amygdala of stressed rats, and females showed markedly higher IL1β as compared to males. Light-dark test (**E**) and open field test (**F**) were performed to assess anxious behavior. Female rats exhibited significantly pronounced anxiety levels after stress than their male counterparts, as evident by their reluctance to spend time in the light chamber (higher dark/light ratio) of the light-dark test and their hesitation to explore the central area of the open field test. Y-maze test (**G**) and novel object recognition (**H**) were performed to assess working memory. Physical stress significantly enhanced the working memory deficits in male rats as compared to females, as depicted by their significantly lower entries to the novel arm in the Y-maze, and less time spent with the novel object in the novel object recognition test. All data are presented as mean with 95% CI (n=10-12); p-values are indicated for specific pairwise comparisons.

Next, we investigated IL1β levels in three areas of the brain that are implicated in anxiety, depression and stress in literature, i.e. the hippocampus, the amygdala, and the prefrontal cortex (Figure 1C). The results were analyzed by three separate 2X2 ANOVAs with stress condition (control, stress) and sex (males, females) as factors. While control levels of IL1β across all three areas of the brain stayed consistent and similar, both psychosocial and physical stress elevated IL1β levels in these brain areas. This upregulation was more robust in the hippocampus and amygdala, especially in females. In the hippocampus, the ANOVA revealed a significant interaction effect between stress type and sex [F(2, 64) = 9.53, p < 0.001, ηp² = 0.229], with females showing increased IL1β in the physical stress model [compared to stressed males, M difference = 146.08, SE = 26.5, t(64) = 5.51, p < 0.001], as well as in the psychosocial stress model [compared to stressed males, M difference = 141.86, SE = 25.9, t(64) = 5.47, p < 0.001]. Similar observations were found in the amygdala, with a significant interaction between stress and sex [F(2, 64) = 7.71, p < 0.001, ηp² = 0.194], and a more robust upregulation in females both in physical [compared to stressed males, M difference = 120.9, SE = 30.9, t(64) = 3.91, p = 0.003] and psychosocial stress [compared to stressed males, M difference = 184.0, SE = 30.2, t(64) = 6.09, p < 0.001] models. In both the hippocampus and amygdala, there were starkly significant differences between control and stressed conditions. In the prefrontal cortex, stress conditions did lead to elevated IL1β levels; however, to a lesser extent than in the limbic structures [F(2, 64) = 14.54, p < .001, ηp² = 0.312], and neither sex nor interaction term was significant. IL1β levels increased in animals undergoing physical stress compared to control animals [females: M difference = −147.9, SE = 36.0, t(64) = −4.11, p = 0.002; males: M difference = −100.9, SE = 34.5, t(64) = −2.93, p = 0.071], as well as psychosocial stress, but only in females [M difference = −135.2, SE = 35.2, t(64) = −3.84, p = 0.004]. Analysis of IL1β in CSF also showed similar trends and significant upregulation of IL1β in both models of stress (Figure 1D). Notably, there was no significant difference in the pattern of upregulation of IL1β between the two stress models, except in the prefrontal cortex, as was the case in plasma and CSF. Additionally, we also characterized the elevation of cortisol in control and stressed animals across both models of stress (Supplementary Figures 1A and 1B) at various time points during the stress regime. These were analyzed with a 2X2X8 mixed factorial ANOVA with stress, sex, and time as factors. Stressed females consistently displayed elevated cortisol levels compared to males, and stressed animals overall displayed peaks in plasma cortisol consistent with extant literature.

Next, we characterized the behavioral phenotype in the models of stress (Figures 1E-1H). We utilized well-validated tests of anxious behavior in rodents, i.e., EPM, OFT, and LDT. We also utilized NOR and Y-maze tests to assess working memory, which is known to be impaired in stress. The methods section outlines the behavior setup and detailed protocols for each of these tests. In brief, to analyze for the LDT, rats were placed in a chamber with a well-lit area with an opening to the dark side of the chamber for 10 min. The ratio of time spent in the dark side versus the light half of the test chamber (D/L ratio) provides a measure of the rat’s anxiety. The higher D/L ratios indicate greater anxiety in the animal. In control animals, the time spent in both chambers was nearly similar, though rats exhibited a marginal preference for the dark side or arena, which is statistically insignificant. Both stress paradigms induced elevated D/L ratios in animals subjected to physical and psychosocial stress. However, this ratio was particularly high in female animals in both models of stress. Statistically, a 2 (male, female) X 3 (control, physical stress, psychosocial stress) factorial ANOVA demonstrated significant main effects of stress type [F(2, 64) = 62.00, p < 0.001, ηp² = 0.660] and sex [F(1, 64) = 55.20, p < 0.001, ηp² = 0.463], as well as a significant interaction between stress and sex [F(2, 64) = 15.00, p < 0.001, ηp² = 0.319], which was evaluated by simple main effects analysis. Under physical stress, females had a significantly higher D/L ratio compared to males [M difference = −1.99, SE = 0.33, t(64) = −6.02, p < 0.001]. Additionally, females exhibited a significantly higher D/L ratio compared to the control group [M difference = −3.31, SE = 0.34, t(64) = −9.79, p < 0.001]. Males also showed increased D/L ratio compared to the control group [M difference = −1.26, SE = 0.32, t(64) = −3.90, p = 0.003]. Under psychosocial stress, females also exhibited a higher D/L ratio compared to males [M difference = −2.29, SE = 0.32, t(64) = −7.07, p < 0.001]. Additionally, females exhibited a significantly higher D/L ratios compared to the control group [M difference = −3.38, SE = 0.33, t(64) = −10.22, p < 0.001], while males also showed an increased D/L ratio compared to the control group [M difference = −1.04, SE = 0.32, t(64) = −3.21, p = 0.031]. The males and females in the control group showed no difference in their D/L ratio.

This pattern was with similar effect sizes and post hoc t scores in OFT, wherein we quantified the time spent by rats in central region of the open field arena (Figure 1F). Anxious rats spent less time in the central area of the arena and spent more time trying to find an escape route along the periphery of the arena. In contrast, the control animals explored the central area as well. This difference is clearly reflected in our analysis. In both physical and psychosocial paradigms, time spent in the central arena fell sharply, and the decline was more pronounced and significant in females. The trends in the EPM were also similar (Figure 2G). In this test, the rats were placed in a 100 cm high plus-shaped maze with two arms without the side wall called open arms that induced fear from height. When rats are placed at the center of the maze and allowed to explore for 5 min, naïve rats display a preference for spending more time in the closed vs. open arms. However, anxiety is associated with a further sharp decline in the time spent in open arms. Our results faithfully replicated this pattern (Figure 2G). In both models of stress, the time spent in the open arm of the EPM decreased significantly, and females showed a more robust decline than males.

**Figure 2.**
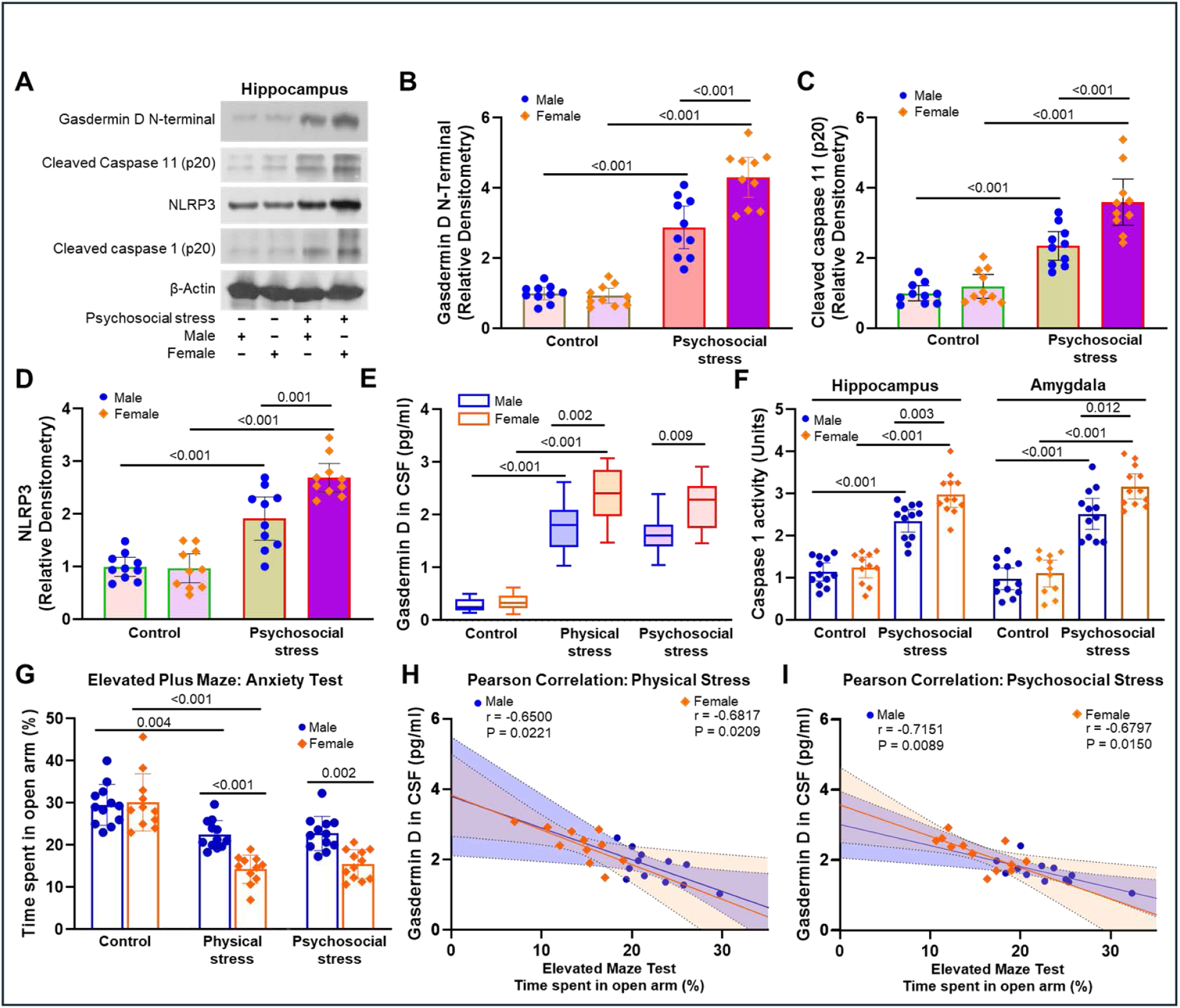
Psychosocial stress induces both canonical and non-canonical NLRP3 inflammasome pathways. **(A)** Immunoblotting was performed to analyze protein levels of gasdermin D N-terminal, cleaved caspase 11, NLRP3, cleaved caspase 1, and β-actin from hippocampal homogenates of male and female rats with (+) or without (-) psychosocial stress. Densitometric analysis showed significantly higher levels of gasdermin D N-terminal **(B)**, cleaved caspase 11 **(C)**, and NLRP3 **(D)** under psychosocial stress. Gasdermin D level **(E)** in the CSF were assessed by ELISA assay, and caspase 1 activity in brain homogenates was measured by activity assay kit **(F)**. Higher levels of cleaved caspase 1 and NLRP3 levels under psychosocial stress indicated induction of the canonical inflammasome pathway in the brain. Enhanced levels of cleaved caspase 11 and gasdermin D N-terminal suggest activation of the non-canonical inflammasome pathway in the brain, which was significantly higher in females than in male counterparts. Elevated plus maze **(G)** was performed to assess anxiety-like behaviors. Female rats under physical and psychosocial stress showed heightened anxiety-like behavior than their male counterparts, as they spent significantly less time in the open arms of the elevated maze. Elevated plus maze scores and gasdermin D levels in CSF under physical stress **(H)** and psychosocial stress **(I)** were subjected to Pearson correlation analysis. Both male and female rats have shown moderate to high negative correlation with gasdermin D levels in CSF and time spent in open arms (less time in open arms represents more anxiety). All data are presented as mean with 95% CI (n=10-12); p-values are indicated for specific pairwise comparisons.

Next, we investigated working memory in stressed vs. control rats. We performed a Y-maze test as the first assessment tool for working memory, where we tested spatial memory. Briefly, the test rat was allowed to explore the Y-shaped testing apparatus with one blocked arm. During the next test trial, this block was removed, and the rat was let free to explore in any direction. Control rats usually display a preference for the newly unblocked arm of the Y-maze to explore. However, stress is usually associated with an abolishment of this preference and, by extension, indicates an impairment in working memory. Our analysis also revealed that both physical and psychosocial stress were associated with a fall in the percentage entry into the novel arm compared to controls (Figure 1G).

Statistical analysis with a 2X3 factorial ANOVA revealed significant main effects of stress and sex type as well as significant interactions [F(2, 64) = 3.21, p = 0.047, ηp² = 0.091], which was further analyzed by a simple main effect analysis. Both physical stress and psychosocial stress led to a reduction in open arm entries. However, interestingly, this difference was more pronounced in male animals, particularly after physical stress [compared to controls: M difference = 19.96, SE = 3.68, t(64) = 5.418, p < 0.001; compared to stressed females: M difference = 12.05, SE = 3.77, t(64) = 3.20, p = 0.032]. In the psychosocial stress condition, both females [M difference = 13.87, SE = 3.77, t(64) = 3.68, p = 0.007] and males [M difference = 18.62, SE = 3.68, t(64) = 5.05, p < 0.001] showed significantly lower entries compared to the control group. We validated these results with the NOR task, which is also a well-accepted behavioral test for working memory in rodents. In this test, rats were first allowed to explore two identical objects in a fixed location in an arena. After a short time, they were reintroduced into the arena, however, with one of the objects replaced with a non-identical novel object. Control unstressed rats with intact working memory display a preference for the novel object by spending more time around it, exploring it by sniffing and poking it with their snouts. A fall in this preference indicated by the time they spent around the novel object denoted a decline in working memory. Our analysis of NOR showed a pattern similar to the Y-maze test (Figure 1H).

Control rats, both male and female, preferentially spent time with the novel object. However, time spent with the novel object declined in stressed animals, signified by an interaction between stress and sex [F(2, 64) = 13.4, p < .001, ηp² = 0.295]. This test also replicated the phenomenon that stress induces significant working memory deficits in males, as compared to females. Under the physical stress condition, males spent significantly less percent time with the novel object compared to the control group [M difference = 16.66, SE = 2.44, t(64) = 6.83, p < 0.001], while females showed no significant difference [M difference = 1.84, SE = 2.55, t(64) = 0.72, p = 1.000]. Similarly, in the psychosocial stress condition, males spent significantly less time with the novel object compared to the control group [M difference = 14.65, SE = 2.44, t (64) = 6.01, p < 0.001], but females again showed no significant difference [M difference = −1.82, SE = 2.49, t(64) = −0.73, p = 1.000].

Notably, in the behavioral tests as well as in IL1β levels in the plasma, brain areas and the CSF, both stress models did not significantly differ in their ability to elicit a phenotype, although both demonstrated sexually divergent patterns of the respective phenotypes.

### Both psychosocial and physical stress models displayed a sexually divergent upregulation of canonical and non-canonical mediators of the inflammasome pathway

We next investigated the levels of the non-canonical (gasdermin D and caspase 11) and canonical (NLRP3) mediators of the inflammasome pathway. Caspase 1 participates in both canonical and non-canonical inflammasome pathways. Immunoblot analysis of the hippocampal lysates from psychosocially stressed rats demonstrated upregulated GSDMD N-terminal, which is produced by cleavage of GSDMD by caspase 1 or caspase 11 (Figure 2A). Immunoblots also showed prominent upregulation of NLRP3 and p20 fragment of cleaved caspases 1 and caspase 11, suggesting their activation. Relative densitometric measurement of band intensities normalized against β-Actin showed significantly increased band intensities for GSDMD-N (Figure 2B), p20 fragment of cleaved caspase 11 (Figure 2C), NLRP3 (Figure 2D), and p20 fragment of cleaved caspase 1 (Supplementary Figure 1C) in the hippocampal lysates of rats subjected to psychosocial stress compared to controls, particularly in females. Data were analyzed by a 2 (stress, control) X 2 (male, female) ANOVA. Interestingly, we found both physical and psychosocial stress resulted in an increased amount of GSDMD in CSF, and this increase was more prominent in female rats (Figure 2E). We further investigated caspase 1 enzyme activity in the hippocampus and amygdala of rats under both physical (Supplementary Figure 1F) and psychosocial stress (Figure 2F). Under both stress paradigms, caspase 1 activity rose in both the hippocampus and amygdala, with female animals showing higher levels in both groups compared to control rats. In control rats, there were no male/female differences in either GSDMD levels or caspase 1 activity.

We also calculated Pearson’s correlation coefficients between the rats’ performance on two behavioral assays, namely the percentage time spent in the open arms of the EPM (Figure 2H-I) and the time spent in the central part of the OFT arena (Supplementary Figure 1D-E) with the GSDMD levels in the CSF. The levels of GSDMD in CSF correlated negatively with the percentage of time spent in the open arms of the EPM in both males (r = –0.650, p = 0.022, df = 10) and females (r = –0.682, p = 0.021, df = 9) under physical stress (Figure 2H). Similarly, under the psychosocial stress condition, there was a significant negative correlation between GSDMD levels in CSF and the percent time spent in the open arm of the EPM in both males (r = –0.715, p = 0.009, df =10) and females (r = –0.6797, p = 0.015, df =10; Figure 2I). We also calculated correlations between CSF levels of GSDMD and time spent in the central square of the OFT arena (Supplementary Figure 1D-E). The analysis revealed a significant negative correlation between open arm time and GSDMD levels in CSF in males (r = – 0.694, p = 0.012, df = 10) and females (r = –0.633, p = 0.037, df = 9) under physical stress (Supplementary Figure 1D). Similarly, under psychosocial stress (Supplementary Figure 1E), we also found significant negative correlations between GSDMD levels in CSF and the time spent in the central square of the OFT arena in males (r = –0.708, p = 0.010, df = 10) and females (r = –0.663, p = 0.019, df = 10). The correlation data suggest that elevated GSDMD levels in the CSF correlated with a higher degree of anxious behavior in both of our stress models.

Until this point, we have provided a thorough behavioral characterization of both stress models that we developed, using markers of both neuroimmune (systemic and brain) and behavioral (anxiety, working memory). We also established that both canonical and non-canonical mediators of the inflammasome pathway were upregulated in stress models. We showed that enhanced levels of GSDMD in CSF are associated with hyper-anxious behavior. Notably, analysis of most of these markers indicated no significant difference between physical stress and psychosocial stress models. The psychosocial stress model, which was established with a repeated social defeat paradigm, produced a similar neuroimmune and behavioral phenotype post-stress. Because the psychosocial stress paradigm is conceptually closer to psychological stress experienced by humans, we conducted the next series of pharmacological experiments in the model of psychosocial stress. Henceforth we refer to this as ‘stress’ for the rest of this article.

### Pharmacological inhibition of caspase 11 by wedelolactone reduces neuroinflammation, improves anxiety, working and fear memory, and mitigates fear acquisition

We utilized wedelolactone, an established caspase 11 inhibitor, to determine the role of the non-canonical inflammasome pathway in mediating anxiogenesis. Figure 3A shows the design of the experiment with drug dosage, route of administration, and timeline. Detailed protocols have been described in the methods section. We measured IL1β levels in the hippocampal lysates with an ELISA assay (Figure 3B). A 2 (control, stress) X 2 (male, female) X 2 (vehicle, drug) factorial ANOVA was utilized to analyze the results. The results showed significant main effects for stress, sex, and drug treatment on hippocampal IL1β levels. Additionally, significant two-way interactions were observed between stress and sex [F(1, 74) = 22.79, p < 0.001, ηp² = 0.235] and between stress and drug [F(1, 74) = 29.29, p < 0.001, ηp² = 0.284]. The interaction between stress and drug and the three-way interaction term were not significant. Simple main effect analysis on the two significant interaction terms was conducted. Post hoc pairwise comparisons with Bonferroni correction for stress and sex factors showed that under the psychosocial stress condition, females exhibited significantly higher IL1β levels compared to the control group [M difference = −335.84, SE = 22.9, t(74) = −14.70, p < 0.001], as did males [M difference = −181.58 SE = 22.9, t(74) = −7.95, p < 0.001]. Comparisons between sexes within the psychosocial stress condition revealed that females had significantly higher IL1β levels than males [M difference = 156.91, SE = 22.0, t(74) = 7.14, p < 0.001]. Bonferroni-corrected pairwise comparisons for stress and drug treatment revealed no significant difference between vehicle and wedelolactone-treated groups in the control condition [M difference = –22.9, SE = 23.7, t(74) = –0.97, p = 1.000]. Under the psychosocial stress condition, vehicle-treated stressed animals exhibited significantly higher IL1β levels compared to the control group [M difference = −346.2, SE = 23.2, t(74) = −14.94, p < 0.001]. Similarly, wedelolactone-treated animals under psychosocial stress also showed higher IL1β levels compared to the control group [M difference = −171.3, SE = 22.5, t(74) = −7.60, p < 0.001]. Within the psychosocial stress condition, however, animals treated with wedelolactone had significantly lower IL1β levels compared to vehicle-treated animals [M difference = 152.0, SE = 22.0, t(74) = 6.92, p < 0.001)]. A very similar pattern was observed in IL1β levels measured in amygdala lysates (Supplementary Figure 2A) and CSF assayed with ELISA (Figure 3C). Once again, wedelolactone reduced the elevated levels of IL1β in stressed animals compared to vehicle-treated animals. The elevation and the pharmacological effect of IL1β were greater in females. Similar trends were found in caspase 1 activity assays in lysates from the hippocampus (Figure 3D) and the amygdala (Supplementary Figure 2B). GSDMD levels in CSF also followed the same pattern. Wedelolactone brought down the GSDMD levels in stressed rats compared to vehicle-treated rats (Supplementary Figure 2C). We then proceeded to measure behavioral outcomes following inhibition of caspase 11 in stressed and control rats with wedelolactone vs. vehicle treatment. In measures of anxious behavior, viz., LDT (Figure 3E) and OFT (Figure 3F), we saw the same pattern of heightened anxiety in stressed rats, particularly in females, and this was significantly mitigated in wedelolactone-treated stressed rats as compared to vehicle-treated stressed rats. Working memory was assessed with the Y-maze test (Figure 3G) and the NOR task (Supplementary Figure 2D). Once again, the same patterns were seen in stressed vs. control animals as were shown in Figures 1. Working memory deficits were seen in stressed animals; however, more in males than females. This was significantly mitigated by wedelolactone treatment.

**Figure 3.**
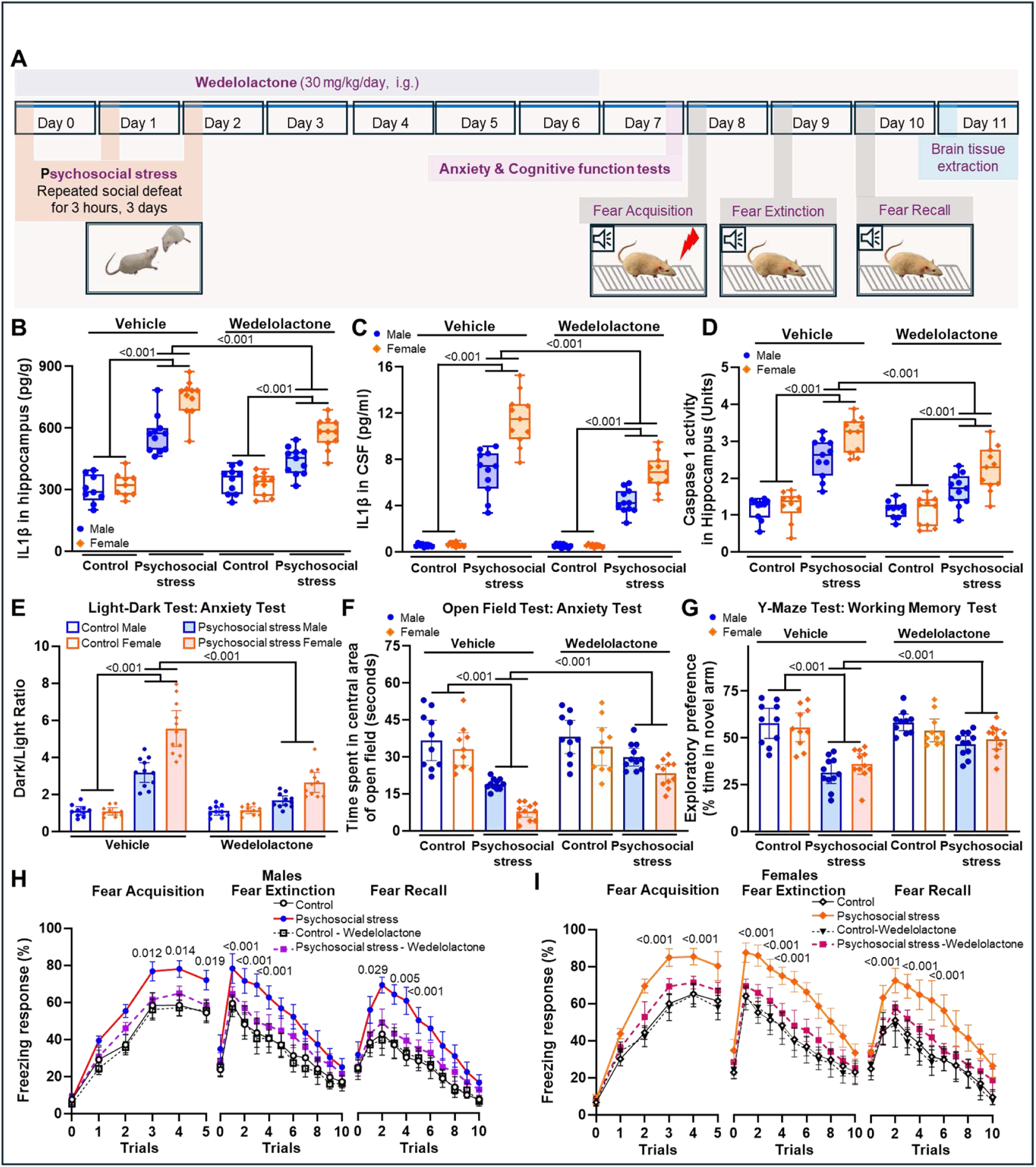
Wedelolactone attenuates IL1β and behavioral dysfunction in rats with psychosocial stress. **(A)** SD rats subjected to repeated social defeat-induced psychosocial stress were dosed with wedelolactone, a caspase 11 inhibitor, or vehicle for 7 consecutive days, starting from the first day of stress induction. IL1β levels in hippocampal homogenates **(B)** and CSF **(C)** were measured by ELISA assay. Wedelolactone treatment significantly reduced the IL1β levels in rats with psychosocial stress. IL1β is processed by activated caspase 1, and to examine its activity after wedelolactone treatment, we performed a caspase 1 activity assay **(D)**. Wedelolactone treatment significantly inhibited caspase 1 activity in rats subjected to psychosocial stress. We performed a light-dark test **(E)** and an open field test **(F)** to evaluate anxiety-like behavior following wedelolactone treatment. Wedelolactone-treated stressed rats spent significantly more time in the light chamber of the light-dark test and in the central area in the open field test than vehicle-dosed stressed group, suggesting the anxiolytic effect of wedelolactone. Working memory function also improved following wedelolactone treatment, as evident by significantly higher preference for novel arms than the vehicle group in the Y-maze test **(G)**. Anxiety is associated with heightened fear memory responses such as enhanced fear acquisition, extinction, and recall. To determine the fear memory response, the fear acquisition, extinction, and recall experiments were performed over 3 days in male **(H)** and female rats **(I)**. Wedelolactone treatment significantly attenuated the fear memory responses in male and female stressed rats. All data are presented as mean with 95% CI (n=10-12); p-values are indicated for specific pairwise comparisons.

Next, we conducted a fear conditioning paradigm to test different aspects of fear memory in these animals (Figures 3H and 3I). Animals were fear conditioned on day 1 (day 8 of the timeline in Figure 3A), and extinction and recall were tested on the second and third consecutive days (days 9 and 10 of the timeline shown in Figure 3A), respectively. A detailed protocol has been explained in the methods section. Briefly, the animals acquired a fear response to a sound presented for 20 sec (conditioned stimulus) coupled with a brief foot shock (unconditioned stimulus) that was delivered through the metal floor grid. The percentage of time they spent freezing during the conditioned stimulus was measured as an indicator of fear acquisition. All rats acquired fear within 3-5 trials; however, both male and female vehicle-treated stressed animals displayed a higher rate of freezing, (Figures 3H and 3I). Wedelolactone treatment significantly reduced the percentage of freezing responses in stressed animals. Through both days of extinction and recall testing, when the sound (conditioned stimulus) was presented to the rats without the noxious foot shock stimulus, wedelolactone-treated animals demonstrated reduced freezing as compared to vehicle-treated rats. We analyzed this data with a 2 (stress, control) X 2 (drug, vehicle) X 3 (time) mixed factorial ANOVA with 2-3 trials per day, as indicated by p values in Figures 3H and 3I. However, even from the plots in Figures 3H and 3I graphed with mean and 95% CIs, it is evident that on the acquisition day vehicle-treated stressed animals started displaying mean freezing responses with non-overlapping CIs from the 3rd trial onwards compared to all other groups, including the wedelolactone-treated male and female stressed rats. Likewise, on days 9 and 10, when extinction and recall were tested, the vehicle-treated stressed rats continued to display higher freezing responses with CIs that did not overlap with any other groups for most of the trials in both males and females, though the traces in female rats showed a better separation. Taken together, this data suggests that inhibition of caspase 11 with wedelolactone treatment attenuated IL1β levels in the brain and improved behavioral outcomes in stressed rats.

### Pharmacological inhibition of GSDMD by disulfiram reduces IL1β and improves behavioral function in stressed rats

We used disulfiram to allosterically inhibit GSDMD cleavage mediated by caspases 1 and 11. Disulfiram is an FDA-approved drug that is widely used in treating alcohol use disorder. The experimental timeline and treatment regimen, route of administration, and design are shown in Figure 4A. Plasma, area-specific brain lysates, and CSF from vehicle and drug-treated control and stressed rats were assessed for levels of IL1β with ELISA assays (Figures 4B-D). The data were analyzed with a 2 (stress, control) X drug (disulfiram, vehicle) X sex (male, female) factorial ANOVA. IL1β level in plasma was elevated in stressed rats, with more pronounced induction of IL1β in females. Disulfiram treatment did not alter levels of IL1β in the plasma in control animals; however, it reduced significantly in stressed rats compared to vehicle-treated groups (Figure 4B). However, the treatment did not mitigate the elevated IL1β levels comparable to control levels. Similar results were obtained for IL1β levels in the hippocampus and amygdala (Figure 4C). To validate our results with disulfiram and confirm that these findings came from the inhibition of GSDMD cleavage, we also treated a third group of rats with necrosulfonamide, a potent inhibitor of necrosis and pyroptosis mediated by GSDMD [73]. Treatment with necrosulfonamide also reduced IL1β levels in the CSF, in a pattern similar to disulfiram treatment (Figure 4D). When we tested these animals for behavioral functions, the disulfiram-treated rats displayed statistically significant reductions in anxiety. Exemplar traces from the OFT arena show a marked reduction of central square crossing in vehicle-treated stressed female rats, compared to controls (Figure 4E). Stressed female rats treated with disulfiram showed some restoration in central square crossings, indicating attenuation of anxiety-like behavior. This is reflected in the quantification of the data in Figure 4F. The data were analyzed with a 2X2X2 factorial ANOVA, as described above. Apart from main effects, significant two-way interactions were observed between stress and sex [F(1, 84) = 9.61, p = .003, ηp² = 0.103] and between stress and drug [F(1, 84) = 20.39, p < .001, ηp² = 0.195]. Post hoc pairwise comparisons with Bonferroni correction for stress and sex factors revealed that under the psychosocial stress condition, females spent significantly less time in the central arena compared to the control group [M difference = 23.58, SE = 2.66, t(84) = 8.87, p < .001], as did males [M difference = 11.92, SE = 2.66, t(84) = 4.49, p < .001].

**Figure 4.**
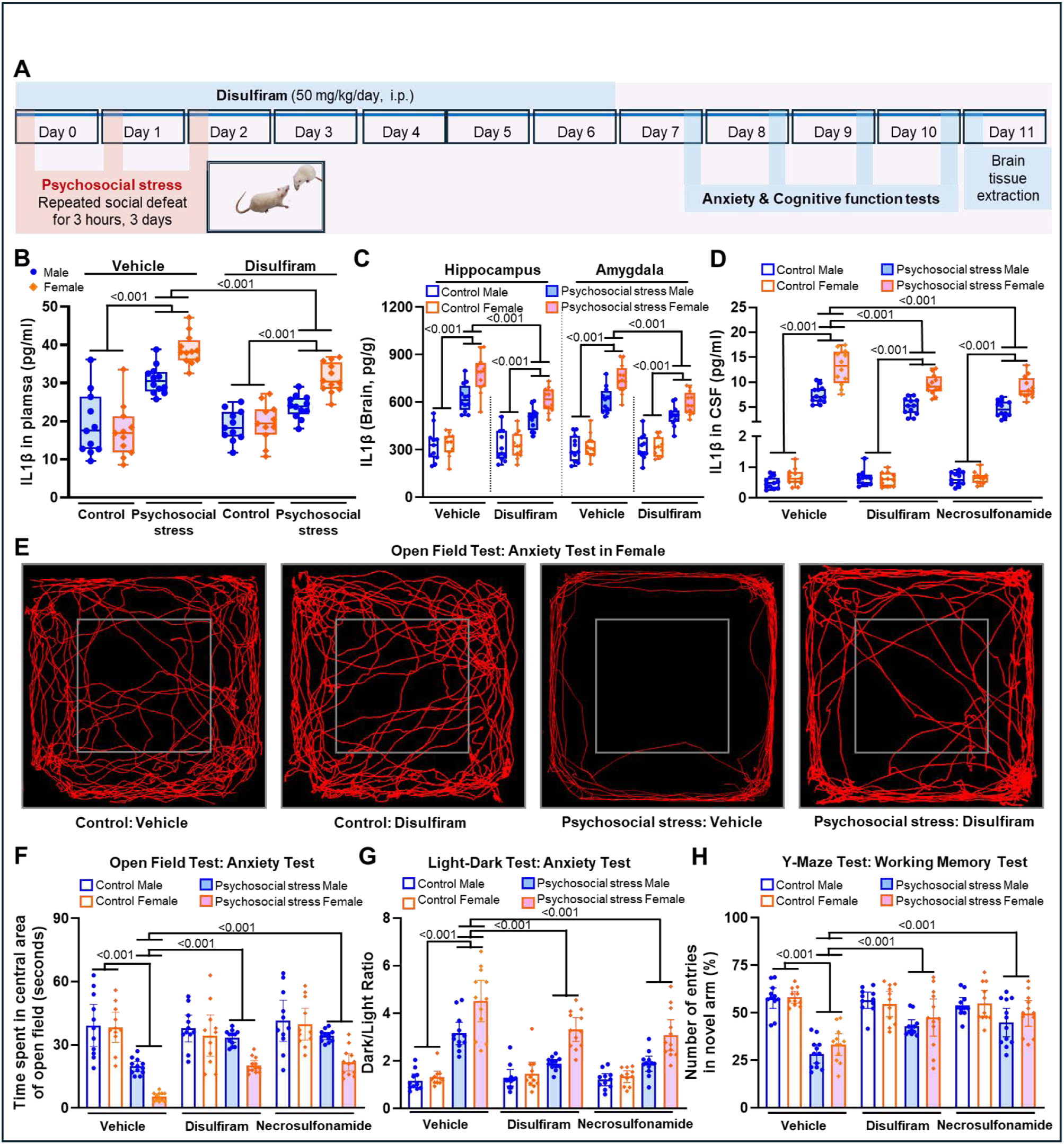
Disulfiram, a gasdermin D inhibitor, suppresses IL1β and anxious behavior following repeated social defeat. **(A)** SD rats subjected to repeated social defeat-induced psychosocial stress received disulfiram or vehicle for 7 consecutive days starting from the first day of stress induction. IL1β levels in plasma **(B)**, hippocampal and amygdala homogenates **(C),** and CSF **(D)** were measured by ELISA assay. Disulfiram treatment significantly reduced the IL1β levels in rats subjected to repeated social defeat. To validate the observations from disulfiram studies, another gasdermin D inhibitor necrosulfonamide was also used **(D)**. Necrosulfonamide treatment significantly reduced IL1β levels in CSF following psychosocial stress, suggesting contribution of gasdermin D to the induction of IL1β levels in the brain. To evaluate the effect of disulfiram treatment on anxiety-like behavior, the open field test **(E-F)** and the light-dark test **(G)** were performed. Animals with anxiety prefer to stay close to the walls of the open field than the central open area of the box during the open field test. Disulfiram and necrosulfonamide treatment significantly reduced the anxiety levels following repeated social defeat, as evident by the higher time spent in the central area of the open field test and more time spent in the light chamber of the light-dark test. Disulfiram and necrosulfonamide treatment also improved working memory in stressed rats, as observed by a higher number of entries in the novel arm of the Y-maze test **(H)**. All data are presented as mean with 95% CI (n=10-12); p-values are indicated for specific pairwise comparisons.

Comparisons between sexes within the psychosocial stress condition revealed that females spent significantly less time in the central arena than males [M difference = –13.79, SE = 2.60, t(84) = – 5.31, p < .001]. Under the psychosocial stress condition, vehicle-treated animals spent significantly less time in the central arena compared to the control group [M difference = 26.23, SE = 2.66, t(84) = 9.87, p < .001], as did disulfiram-treated animals [M difference = 9.26, SE = 2.66, t(84) = 3.48, p = .005]. Within the psychosocial stress condition, however, disulfiram-treated animals spent significantly more time in the central arena compared to vehicle-treated animals [M difference = 14.29, SE = 2.60, t(84) = 5.50, p < .001]. The necrosulfonamide-treated group of rats also showed the same pattern of reduction of anxious behavior in stressed animals as compared to vehicle-treated ones. Treatment of stressed rats with disulfiram or necrosulfonamide improved working memory function, as indicated by the higher time spent in the light area in the LDT (Figure 4G). Y-maze test results further validated that inhibition of GSDMD with disulfiram and necrosulfonamide mitigated the stress-induced reduction in working memory, which was especially more prominent in males (Figure 4H).

### A combination treatment with pharmacological inhibitors of both the canonical and non-canonical mediators of the inflammasome pathways offers better behavioral rescue, mitigation of spine density reduction, and more robust amelioration of neuroinflammation in stressed rats

In this phase, we investigated the potential benefit of simultaneous inhibition of canonical and non-canonical inflammasomes by employing a combination therapy approach wherein we used both types of inhibitors and then compared them to individual drug-treated groups and vehicle-treated groups.

The drugs, dosages, administration routes, and timeline of drug administration, induction of stress, and assessment are all outlined in Figure 5A. For the first set of experiments, we performed a proof-of-concept investigation to test if the simultaneous inhibition of the NLRP3 inflammasome and caspase 11 will be beneficial by enhanced reduction of IL1β and anxiety levels. To this end, we treated animals with MCC950, a well-known selective inhibitor of NLRP3 inflammasome assembly, and wedelolactone, a caspase 11 inhibitor. These treatments were tested individually and in combination using rats subjected to psychosocial stress. The combination-drug treatment demonstrated lower IL1β levels in the hippocampus, not only relative to the stressed animals but also compared to the individual drug-treated groups (Figure 5B). Specifically, in the hippocampus, a significant interaction between drug treatment and psychosocial stress was observed [F(3, 76) = 32.8, p < .001, ηp² = 0.564]. Notably, comparisons between treatment groups under the psychosocial stress condition revealed that animals treated with MCC950 in combination with wedelolactone had significantly lower IL1β levels compared to animals treated with either only wedelolactone [M difference = −154.82, SE = 31.2, t(76) = −4.96, p < 0.001] or only MCC950 [M difference = −212.92, SE = 31.2, t(76) = −6.82, p < 0.001]. Working memory assessment with the Y-maze task echoed the similar pattern in female rats. Vehicle-treated female rats displayed a worsening working memory function following stress. The combination treatment with MCC950 along with wedelolactone was more efficacious in improving novel arm entries, i.e., working memory, as compared to the individual treatments (Figure 5C). Aside from the main effects, the 2X2 factorial ANOVA revealed a significant interaction between drug treatment and psychosocial stress [F(3, 76) = 11.8, p < .001, ηp² = 0.317]. Comparisons within the psychosocial stress condition revealed that animals treated with a combination of MCC950 and wedelolactone exhibited significantly higher percentages of entries compared to animals treated with MCC950 alone [M difference = 13.12, SE = 2.94, t(76) = 4.47, p < .001]. This suggests that a combination treatment approach of simultaneous blocking both canonical NLRP3 inflammasome assembly and non-canonical caspase 11 activation produces a more robust benefit than using them in isolation.

**Figure 5.**
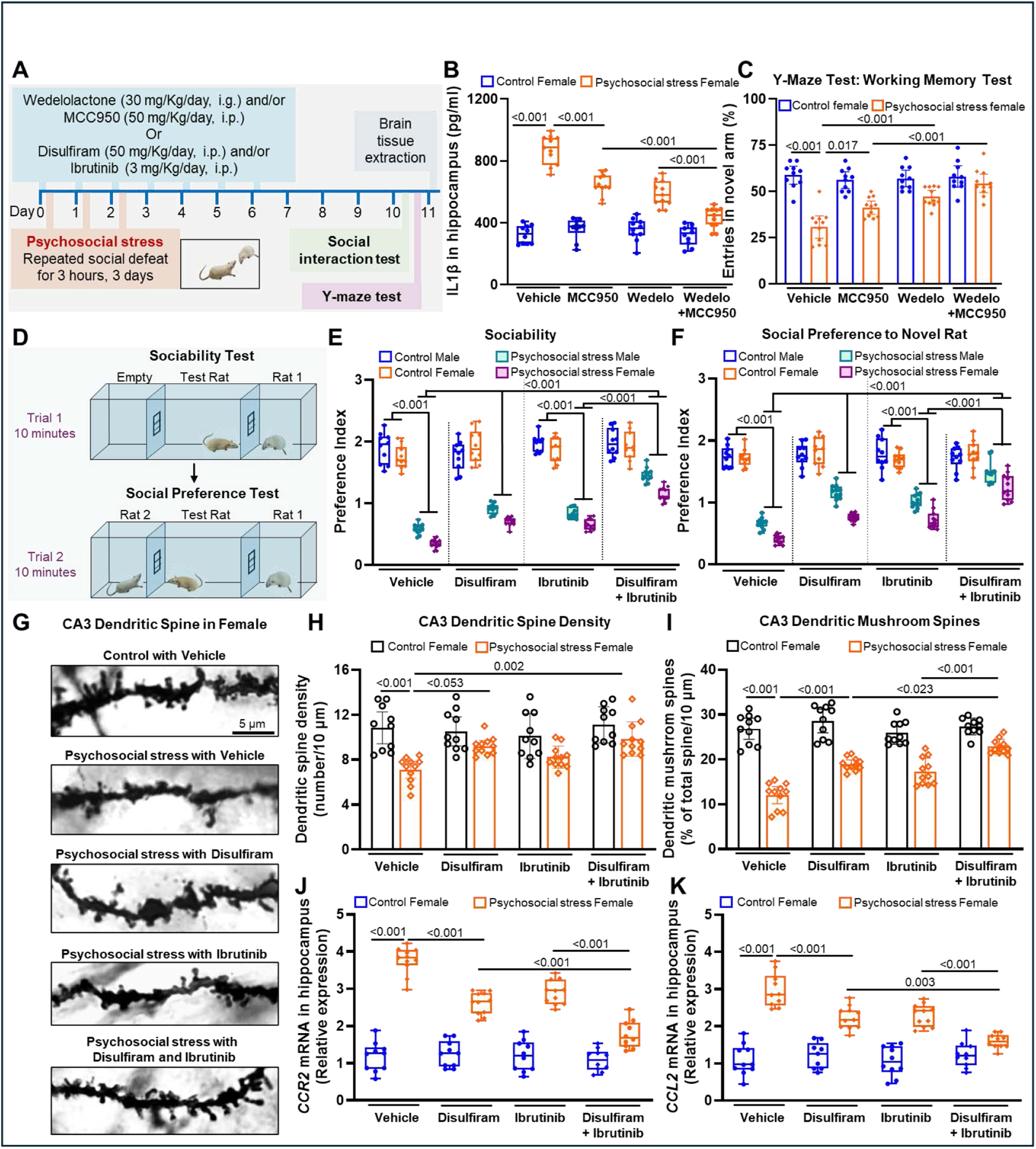
Concurrent inhibition of mediators of canonical and non-canonical inflammasomes potentiates anxiolytic effect. **(A)** Canonical inflammasome pathway was inhibited by MCC950, a specific NLRP3 inhibitor, or ibrutinib, a specific BTK inhibitor. Non-canonical inflammasome pathway was inhibited by wedelolactone, a caspase 11 inhibitor, or disulfiram, a gasdermin D inhibitor. In female rats subjected to repeated social defeat, as compared to individual treatments, a combination treatment with MCC950 and wedelolactone significantly reduced the IL1β levels in the hippocampus **(B)**, and improved working memory as assessed by the Y-maze test **(C)**. We performed social interaction tests to analyze sociability and social preference **(D)**. Sociability index was measured by determining the preference of the test rat to interact with another rat (Rat 1) as compared to an empty chamber. Thereafter, social preference was analyzed by assessing the preference of the test rat to the novel rat (Rat 2) as compared to the familiar rat (Rat 1). As compared to individual treatment of stressed rats, the combinational treatment to inhibit both canonical and non-canonical inflammasomes by simultaneous treatment with disulfiram and ibrutinib was more effective to significantly improve sociability **(E)** and social preference indexes **(F)**. Repeated social defeat indued anxious behavior is associated with downregulation of dendritic spine density and reduction of mushroom-shaped mature dendritic spines. Golgi-cox staining **(G)** was performed to analyze dendritic spine density **(H)** and the number of mushroom-shaped spines **(I)** in the CA3 region of the hippocampus. Following repeated social defeat in female rats, a combination treatment with disulfiram and ibrutinib significantly attenuated the reduction of spine density and the number of mushroom-shaped mature spines. Similarly, the combination treatment with disulfiram and ibrutinib significantly reduced the neuroinflammation, as evident by a significant reduction of mRNA levels of monocyte/macrophage receptor *CCR2* **(J)** and microglial ligand *CCL2* **(K)** in the hippocampus of female rats subjected to psychosocial stress. All data are presented as mean with 95% CI (n=10-12); p-values are indicated for specific pairwise comparisons.

Earlier, we had reported how a BTK inhibitor, ibrutinib, which is also FDA-approved for treating certain kinds of human cancers, can reduce IL1β and attenuate stress-induced hyper-anxious behavior in rodents through inhibition of NLRP3 inflammasome. In the present study, we wanted to suppress both IL1β processing and release with FDA-approved ibrutinib and disulfiram by inhibiting NLRP3 inflammasome formation and GSDMD-mediated pore assembly, respectively. To this end, we treated groups of rats with ibrutinib and disulfiram alone and in combination and compared them to vehicle-treated rats under stress and control conditions, as shown in Figure 5A. As an added metric of behavior analysis, we performed social interaction tests to assess the efficacy of this approach (Figure 5D). Briefly, in the first phase to assess sociability, the test rat was placed in the central chamber of the apparatus, while one of the side chambers contained a rat (Rat 1, same biological sex as the experimental rat), and the other chamber remained empty. The transparent wall and opened windows at either side of the central chamber allowed the test rat to see and smell Rat 1 in one of the side chambers. The recording of the interaction time of the test rat with Rat 1 provided a sociability index. In the second phase, to assess social preference, a novel rat (Rat 2, same biological sex as the test rat) was placed in the second empty side chamber, and the interaction time of the test rat with Rat 1 and Rat 2 was compared to determine the social preference index. Sociability (Figure 5E) refers to the sociability preference index, which was determined by dividing the time spent by the test rat interacting with rat 1 to the time spent near the empty chamber. Social preference index (Figure 5F) was calculated by dividing the time spent by the test rat interacting with the novel rat (Rat 2) by the time spent interacting with the familiar rat (Rat 1).

Both sociability and social preference deteriorated drastically following psychosocial stress in vehicle-treated animals (Figures 5E-F). A 2 (control, stress) X 2 (male, female) X 4 (vehicle, ibrutinib, disulfiram, combination treatment) factorial ANOVA revealed significant two-way interactions between stress and sex [F(1, 152) = 15.32, p < .001, ηp² = 0.092] and between stress and drug [F(3, 152) = 33.83, p < .001, ηp² = 0.400], apart from main effects of all three for the sociability index (Figure 5E). The interaction was further broken down by a simple main effects analysis. The general worsening of social interaction was more prominent in females across all treatment groups. Post hoc pairwise comparisons with Bonferroni correction for stress and drug factors revealed that animals under psychosocial stress treated with vehicle exhibited significantly lower sociability scores compared to those treated with disulfiram [M difference = 0.347, SE = 0.0501, t(152) = 6.93, p < .001], ibrutinib [M difference = 0.279, SE = 0.0501, t(152) = 5.57, p < .001], and disulfiram with ibrutinib [M difference = 0.8404, SE = 0.0501, t(152) = 16.78, p < .001]. However, animals treated with both disulfiram and ibrutinib under psychosocial stress had significantly higher preference index scores compared to those treated with only disulfiram [M difference = 0.494, SE = 0.0501, t(152) = 9.85, p < .001] or only ibrutinib [M difference = 0.561, SE = 0.0501, t(152) = 11.21, p = < .001]. Similar results were obtained for social preference for the novel rat (Rat 2) as well (Figure 5F).

Next, we investigated dendritic spine parameters in CA3 hippocampal neurons of these rats in Golgi-Cox-stained thick sections. Spine counts, density, and specific spine morphologies have been associated with neuronal plasticity [92,93]. Both animal and human studies of depression, anxiety, and stress have demonstrated neuroplastic changes in terms of spine density and morphology [94–96].

Consistent with this, we observed a significant reduction of spine density in apical dendrites of CA3 hippocampal neurons of vehicle-treated stressed female rats, as compared to controls (Figures 5G and 5H). A 2X2X4 factorial ANOVA, as designed above, revealed significant main effects for psychosocial stress and drug treatment on the number of dendritic spines in female rats. Additionally, a significant interaction between psychosocial stress and drug treatment was observed [F(3, 76) = 3.01, p = 0.035, ηp² = 0.106]. Post hoc pairwise comparisons with Bonferroni correction for psychosocial stress and drug treatment factors revealed that stressed animals treated with a combination of disulfiram and ibrutinib showed significantly higher spine density compared to vehicle-treated animals [M difference = 2.727, SE = 0.650, t(76) = 4.20, p = 0.002]. Next, we quantified the number of mushroom spines, i.e., mature spines, in the dendritic segments of CA3 hippocampal neurons, which are associated with better memory performance in cognitive tasks [93]. A 2X2X4 factorial ANOVA showed main effects of stress and drug, qualified by a significant interaction [F(3, 76) = 13.7, p < 0.001, ηp² = 0.351]. As clear from Figure 5G and the analysis in Figure 5I, stress caused a sharp reduction of mushroom spines in dendrites. Although not a full rescue, post hoc pairwise comparisons with Bonferroni correction for psychosocial stress and drug treatment factors revealed that under the psychosocial stress condition, animals treated with a combination of disulfiram and ibrutinib had significantly higher mushroom spine percentages as compared to vehicle-treated animals [M difference = 10.835, SE = 1.12, t(76) = 9.66, p < 0.001]. Furthermore, within the psychosocial stress condition, animals treated with a combination of disulfiram and ibrutinib exhibited significantly higher mushroom spine percentages as compared to animals treated with only disulfiram [M difference = 3.914, SE = 1.12, t(76) = 3.49, p = 0.023] or only ibrutinib [M difference = 5.574, SE = 1.12, t(76) = 4.97, p < 0.001].

We also studied other mediators known to drive neuroinflammatory responses under stress. CCL2, expressed by glial and brain endothelial cells, is a chemokine ligand that binds to chemokine receptor CCR2 on microglia and monocytes to facilitate the recruitment of immune cells to the brain during stress [97–99]. In the repeated social defeat paradigm that we utilized in psychosocial stress, upregulation of both CCL2 and CCR2 is noted in the literature [100,101]. Consistent with this, female vehicle-treated stressed rats displayed a marked elevation of *CCL2* and *CCR2* mRNAs (Figures 5J and 5K). The combination treatment mitigated this elevation better than the individual drugs alone. A 2X2X4 factorial ANOVA for *CCR2* mRNA levels showed main effects for both drug and stress with a significant interaction between them [F(3, 75) = 25.9, p < 0.001, ηp² = 0.509]. When evaluated, post hoc pairwise comparisons with Bonferroni correction for psychosocial stress and drug treatment factors revealed that under the psychosocial stress condition, animals simultaneously treated with disulfiram and ibrutinib exhibited significantly lower upregulation of *CCR2* mRNAs, as compared to vehicle-treated animals [M difference = −1.9879, SE = 0.154, t(75) = −12.95, p < 0.001]. Stressed animals treated with a combination of disulfiram and ibrutinib also had significantly attenuated induction of *CCR2* mRNAs, as compared to those treated with either disulfiram alone [M difference = −0.8352, SE = 0.154, t(75) = −5.44, p < .001] or ibrutinib alone [M difference = −1.1243, SE = 0.154, t(75) = −7.32, p < .001]. Very similar results were obtained following the analysis of *CCL2* mRNA (Figure 5K). In addition, we analyzed the hippocampal homogenates using an ELISA assay for two markers of neuroinflammation, i.e., ICAM1 and CXCL2 (Supplementary Figure 3A, B). ICAM1 is an adhesion molecule whose expression on the surface of endothelial and various immune cells is upregulated during stress and neuroinflammation [100–104]. We also observed elevated expression of ICAM1 in hippocampal lysates from vehicle-treated stressed rats. This elevation of ICAM1 was attenuated slightly by individual treatment with ibrutinib or disulfiram, and to a greater extent by the combination treatment (Supplementary Figure 3B). Similar trends were also observed from CXCL2 analysis, another chemokine associated with exacerbated neuroinflammation [105–107]. In the case of both these mediators, the combination treatment was better at checking stress-induced elevations than the two drugs employed by themselves (Supplementary Figure 3A, B).

Taken together, the data in Figure 5 suggested that a combination treatment approach worked better, i.e., when the mediators of the NLRP3 inflammasome and caspase 11-GSDMD pathways were simultaneously targeted. This resulted in efficient attenuation of IL1β processing and release, which led to suppression of neuroinflammation. This was further supported by orthogonal validation from the analysis of dendritic spines, neuroinflammatory mediator, and behavioral data. We therefore conducted our final set of pharmacological inhibition experiments using a combination treatment of disulfiram and ibrutinib.

### A combination of disulfiram and ibrutinib leads to reduced IL1β induction and improved behavioral function in stressed rats

The design and timeline of this experiment are shown in Figure 6A, along with drug dosages and routes of administration. The combination treatment of stressed rodents with disulfiram and ibrutinib reduced IL1β in lysates from the hippocampus and amygdala (Figure 6B), plasma (Supplementary Figure 4A), and CSF (Supplementary Figure 4B), otherwise starkly increased in vehicle-treated stressed rats. The elevation of IL1β and its mitigation by the combination treatment were more pronounced in female rats. A 2 (stress, control) X 2 (male, female) X 2 (vehicle, combination treatment) factorial ANOVA revealed significant main effects for drug treatment and stress on hippocampal IL1β levels, with significant two-way interactions observed between drug treatment and stress [F(1, 82) = 54.47, p < .001, ηp² = 0.399] and between stress and sex [F(1, 82) = 17.25, p < .001, ηp² = 0.174]. Post hoc pairwise comparisons with Bonferroni correction showed that in the vehicle group, animals under psychosocial stress exhibited significantly higher IL1β levels compared to the control group [M difference = −390.9, SE = 24.3, t(82) = −16.07, p < .001]. Comparisons between treatment groups under psychosocial stress revealed that animals treated with a combination of ibrutinib and disulfiram exhibited significantly lower IL1β levels compared to those treated with a vehicle [M difference = −236.2, SE = 23.2, t(82) = −10.18, p < .001]. Similar results were seen in the amygdala, plasma, and CSF for IL1β. We found the similar trend following analysis of caspase 1 activity in both the hippocampus and the amygdala (Figure 6C).

**Figure 6.**
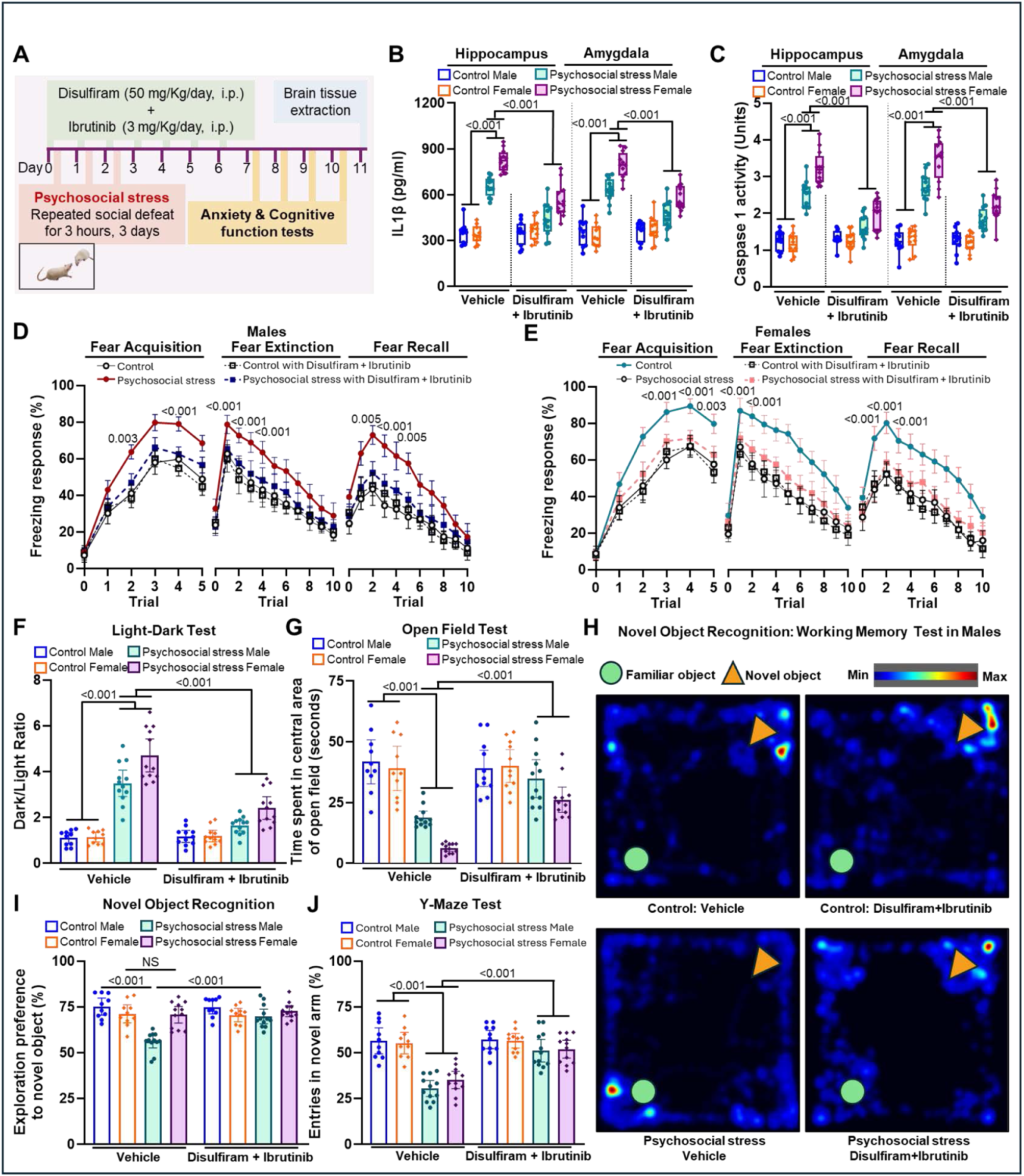
Combination treatment with disulfiram and ibrutinib synergistically improves the anxiolytic effect and working memory. **(A)** A combination treatment with disulfiram and ibrutinib was carried out from the first stress induction day to a total of 7 consecutive days. IL1β levels and caspase 1 activity were measured in the hippocampus and amygdala homogenates of SD rats. The combination treatment significantly reduced the IL1β levels **(B)** and caspase 1 activity **(C)** in the brains of male and female rats subjected to repeated social defeat. This combination treatment also significantly mitigated the anxiogenic fear memory responses, as evident by the reduction of fear acquisition, extinction, and recall responses in male **(D)** and female **(E)** stressed rats. To investigate the effects of combination treatment on anxiety-like behavior, the light-dark test **(F)** and the open field test **(G)** were performed. As compared to individual treatments, a combination treatment significantly reduced anxiety-like behavior, suggesting better anxiolytic potential of the combination treatment. The heat map of the novel object recognition task indicated that stressed rats spent less time with the novel object, as compared to control rats, as depicted by the lesser blue/red color at the top right corner **(H)**. This combination treatment significantly improved working memory function in stressed rats, as evident by the higher time spent with the novel object in the novel recognition object test **(H, I)** and more entries to the novel arm of the Y-maze **(J)**. All data are presented as mean with 95% CI (n=10-12); p-values are indicated for specific pairwise comparisons.

We next assessed fear memory in the animals. Vehicle-treated animals under stress showed higher freezing rates during fear acquisition on day 8, with females showing more freezing (Figures 6D-E). When tested on day 9 for extinction and day 10 for recall, vehicle-treated stressed rats persistently showed higher freezing responses even though they did demonstrate extinction eventually. Notably, in stressed rats the combination treatment with ibrutinib and disulfiram proved more efficient in reducing freezing behavior during fear acquisition, extinction, and recall trials (Figures 6D-E). The behavioral tests for anxiety (LDT and OFT) and working memory (Y-Maze and NOR) also showed significant improvements in the combination therapy group, with vehicle-treated animals doing predictably worse following psychosocial stress (Figures 6F-J). Once again, we observed a higher working memory deficit in male animals, as seen in Figure 6I for the NOR test, where the stressed male rodent spent less time with the novel object. The combination therapy with ibrutinib and disulfiram mitigated the stress-induced working memory deficits, as indicated by the increased exploration time with the novel object in the NOR test and a higher number of entries into the novel arm of the Y-maze (Figures 6I and 6J).

### Cell culture studies demonstrate the role of the NLRP3-caspase 1 and caspase 11-GSDMD pathways in the propagation of inflammatory signaling in PBMCs from stressed rats

In extant literature, the cleavage of GSDMD into its N-terminal is associated with minute pore formation in the cell membrane, which enables the release of IL1β and efflux of K^+^ ions from the cell, potentiating further inflammatory cascades [39]. The influx of K^+^ ions can also trigger activation of both the NLRP3 inflammasome and caspase 1. Further, increase in the number of pore formations in the cell can cause the building of osmotic pressure and an influx of water, followed by cell swelling and eventual lysis. However, this mechanism is yet to be studied in models of psychological stress.

To investigate the non-canonical activation of the caspase 11-GSDMD pathway and its contribution to the release of IL1β and caspase 1 activation, we conducted a set of *in vitro* mechanistic experiments using PBMCs purified from rats under control and stressed conditions. Our prior work with rodents had shown that the activation of the NLRP3 inflammasome pathway under conditions of stress is quite prominent in peripheral immune cells. In our current investigation, we observed that PBMCs purified from stressed animals showed higher levels of GSDMD N-terminal and cleaved caspase 11, indicating the stress-induced activation of non-canonical inflammasomes (Figure 7A).

**Figure 7.**
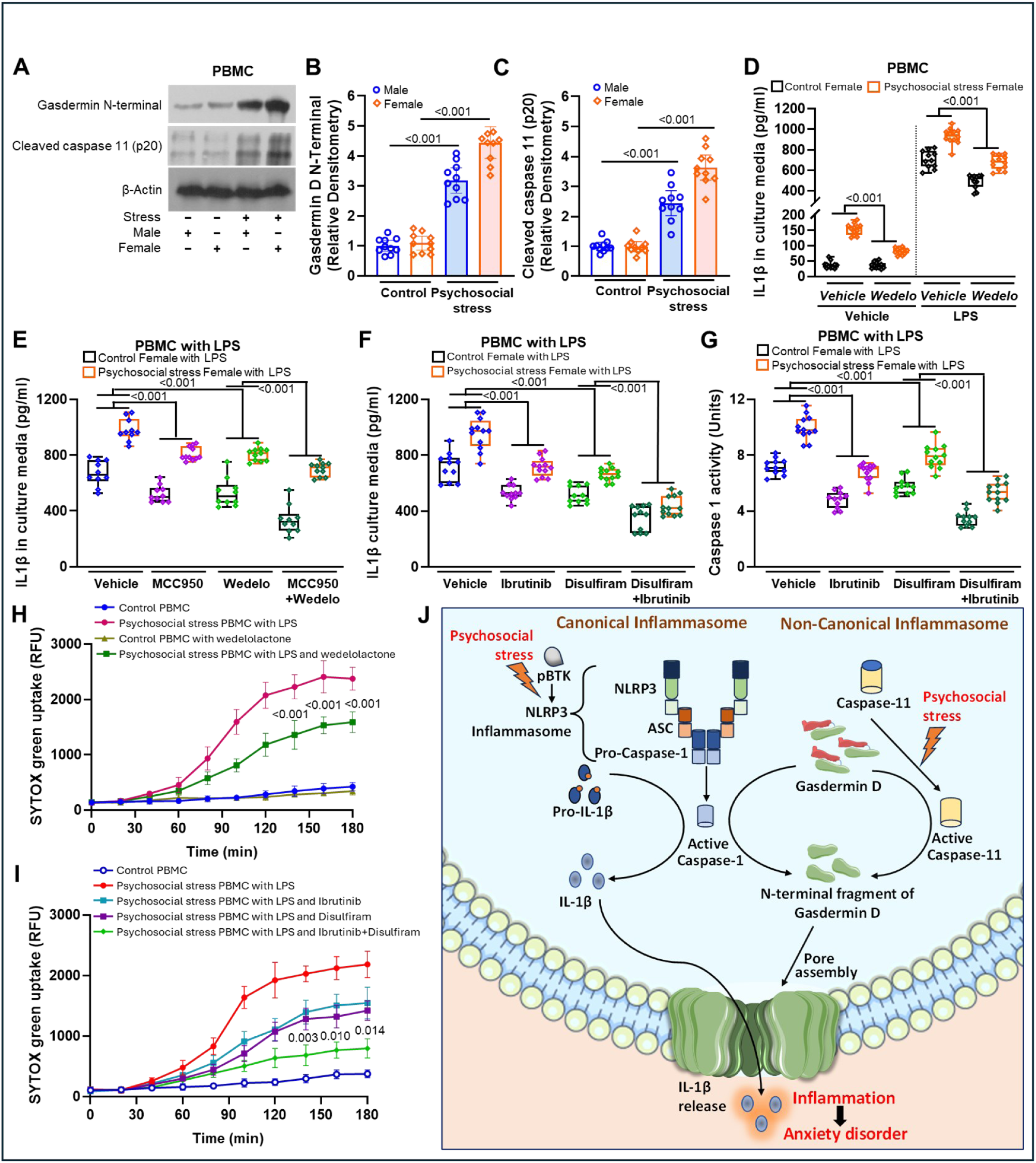
Psychosocial stress induces caspase 11 and gasdermin D, the mediators of the non-canonical inflammasome pathway, in PBMCs. **(A)** Immunoblotting was performed to measure the levels of gasdermin D N-terminal and cleaved caspase 11 from isolated PBMCs of rats with (+) or without (-) psychosocial stress. Densitometric analysis showed significantly higher levels of gasdermin D N-terminal **(B)** and cleaved caspase 11 **(C)** following repeated social defeat. This suggests activation of the non-canonical inflammasome pathway in the peripheral immune cells under psychosocial stress. IL1β levels in the culture media of PBMCs from stressed rats were significantly higher as compared to controls, and wedelolactone treatment significantly reduced IL1β levels **(D)**. A simultaneous inhibition of NLRP3 and caspase 11 by a combination treatment with MCC950 and wedelolactone was able to significantly reduce the IL1β levels in LPS-challenged PBMCs from stressed rats **(E)**. Similarly, concurrent inhibition of BTK and gasdermin D by a combination treatment with ibrutinib and disulfiram significantly reduced IL1β levels in LPS-induced PBMCs from stressed rats **(F)**. IL1β processing enzyme caspase 1 activity was also significantly reduced with ibrutinib and disulfiram treatment **(G)**. To analyze the pyroptosis, the membrane pore formation by gasdermin D, the SYTOX Green uptake assay was performed. Higher pore formation on the plasma membrane causes higher levels of SYTOX Green influx into the cells. The treatment with wedelolactone significantly decreased SYTOX Green uptake in the LPS-induced PBMCs from stressed rats **(H)**. As compared to individual treatment, the combination treatment of disulfiram and ibrutinib was more effective in attenuating SYTOX Green uptake **(I)**. This suggests caspase 11 and gasdermin D are responsible for membrane pore formation. Suggestive schematic diagram of canonical and non-canonical NLRP3 inflammasome pathway in psychosocial stress **(J)**. Artwork adapted from https://smart.servier.com, CC BY 4.0. All data are presented as mean with 95% CI (n=10-12); *p<0.05; one-way ANOVA with Bonferroni post hoc test; p-values are indicated for specific pairwise comparisons.

Even within the isolated cells, the increase was more apparent in cells from female rats under stress (Figure 7A). Relative densitometry of GSDMD N-terminal and cleaved p20 fragment of caspase 11 immunoblots reflect this trend (Figures 7B-C).

To investigate the role of this activation of caspase 11 in the upregulation of IL1β, we treated cultures of PBMCs, isolated from control and stressed rats, with wedelolactone (Figure 7D). A set of PBMCs was primed with lipopolysaccharide (LPS), a component of the bacterial cell walls and known elicitor of inflammatory response. The culture media of naïve/vehicle-treated PBMCs from stressed animals show baseline elevated levels of IL1β to begin with. However, when challenged with LPS, there was a massive upregulation of IL1β in the PBMC cultures. Notably, PBMC cultures from stressed rodents, which were treated with wedelolactone, showed the elevation of IL1β in media was significantly dampened as compared to vehicle-treated PBMCs from stressed rats. Similarly, wedelolactone treatment attenuated the induction of IL1β following LPS challenge. From here onwards, we only show data from LPS-challenged PBMC cultures originating from female animals, which have a more robust neuroinflammatory phenotype, as seen above.

To further investigate the role of canonical and non-canonical inflammasomes in IL1β release from cells, the PBMC cultures from control and stressed rats were treated with respective inhibitors (Figure 7E-F). PBMCs were primed with LPS to elicit a higher inflammatory response. MCC950 and wedelolactone attenuated the release of IL1β from LPS-challenged PBMCs from stressed rats; however, the combination therapy with MCC950 and wedelolactone provided more robust protection from this induction of IL1β (Figure 7E). To further validate this finding, we used another set of inhibitors, i.e., ibrutinib and disulfiram, to block canonical and non-canonical inflammasomes, respectively. Although LPS-challenged PBMC cultures from stressed female rats secreted higher amounts of IL1β in the culture media, this was mitigated significantly in cultures that were treated with ibrutinib, disulfiram, or both (Figure 7F). The mitigation was most prominent in the cultures that received the combination treatment of ibrutinib and disulfiram. Naïve cultures of PBMC from control and stressed animals without LPS challenge also displayed similar findings (Supplementary Figure 5A). The combination treatment with ibrutinib and disulfiram (Figure 7G), as well as MCC950 and wedelolactone (Supplementary Figure 5B), more efficiently attenuated caspase 1 activity in LPS-challenged PBMCs from stressed rats. Together, the studies from PBMCs isolated from stressed rats strongly indicated the critical role of both NLRP3-caspase 1 and caspase 11-GSDMD pathways in the efficient induction of IL1β and the propagation of inflammatory signaling.

Next, to assess the plasma membrane pore formation following the induction of non-canonical GSDMD N-terminal fragments, we performed a time course assay with SYTOX Green, a DNA-binding dye that usually does not permeate across intact cell membranes [89]. As seen in Figure 7H, the PBMC cultures originating from the control animals did not take up the SYTOX Green. However, LPS-challenged PBMC cultures that originate from stressed animals (red trace) steadily take up the SYTOX Green over 180 minutes, supporting the existence of pores in the cell membrane. LPS-challenged cultures from stressed rats treated with wedelolactone, however, take up the dye slower and to a lesser extent (green trace). This suggests that inhibition of caspase 11 in LPS-challenged PBMCs by wedelolactone mitigates the plasma membrane leakage. We repeated this experiment with PBMC cultures with ibrutinib, disulfiram, and a combination of these two drugs (Figure 7I). The PBMC challenged with LPS showed high uptake of SYTOX Green, signifying the pore formation in the plasma membrane. However, cells treated with ibrutinib, disulfiram, and a combination of these two drugs, showed significantly lower SYTOX Green uptake. However, the combination treatment is much better at reducing SYTOX Green uptake (green trace). The p values from the mixed factorial ANOVA are mentioned on the traces of figures 7H and 7I. We also checked whether this leaky membrane led to lytic cell death. To this end, we performed WST-8 cell viability assays [88] on all of the cultures (Supplementary Figure 5C, D). However, we found no evidence of a decline in cell viability in any of these groups.

## Discussion

In this study we built upon the foundation of our earlier work in mice, when we had provided rigorous evidence of the upregulation of the BTK-NLRP3-caspase 1 pathway induced in the models of psychological stress (predator odor) and physical restraint stress. We had also shown that treatment with ibrutinib, an FDA-approved drug for treating certain human cancers, mitigates the neuroinflammation, particularly reducing IL1β and rescuing anxious behavior in stressed rodents [48]. We extend these findings in the present study and investigate canonical inflammasome mediators that facilitate the processing of IL1β and cleavage of GSDMD by caspase 1. We also investigate non-canonical activation of GSDMD by its cleavage by caspase 11. Interestingly, caspase 1 can process pro-IL1β into IL1β as well as cleave GSDMD to its N-terminal and C-terminals [51,108,109]; however, caspase 11 is mainly specific to GSDMD [110,111]. Parallel to our earlier results, several other groups have published results that confirm that the canonical caspase 1-GSDMD pathway is involved in neuroinflammatory responses underlying chronic stress-induced depression as well as a range of other inflammatory diseases [63–65,112–114]. The caspase 11/GSDMD non-canonical inflammasome pathway has been shown to activate during sepsis, lung injury, asthma, skin psoriasis, peptic ulcers, and graft versus host disease; however, the role of non-canonical activation mediated by caspase 11 has not been studied in models of psychosocial stress or psychological disorders. We designed most of the experiments in the present study to fill this gap.

We have provided the first evidence to our knowledge about the role of both canonical and non-canonical inflammasome activation in psychosocial stress. Beside elevated levels of active caspase 1 (p20) and NLRP3 inflammasome, we have also observed elevated levels of active caspase 11 (p20) and GSDMD N-terminal in both the hippocampus and amygdala of the rats subjected to repeated social defeat. This is the first evidence that psychosocial stress can induce caspase 11/GSDMD, the mediators of non-canonical inflammasomes. Both caspase 1 and caspase 11 can initiate GSDMD cleavage and subsequent pore formation. GSDMD is more effectively cleaved by caspase 11, creating membrane pores that cause K^+^ efflux and NEK7 phosphorylation, which in turn activates NLRP3 and creates the NLRP3 inflammasome complex [115]. Thus, our study further implies that psychosocial stress-induced activation of both caspase 1 and caspase 11 can drive induction of NLRP3 inflammasome assembly, which can further activate caspase 1.

We also provide detailed characterization of the rat model of repeated social defeat adapted from the existing literature. This model was further strengthened by placing rats in solitary housing following psychosocial stress induction and then examining different facets of behavior dysfunction. There is evidence that while grouped housing of rodents may mitigate the stress responses, the solitary housing influences corticosterone levels and sleep duration [116]. So, we hypothesized that solitary housing after the stress induction may elicit a more robust response. While characterizing the neuroinflammatory response in stressed animals, we observed a robust elevation of IL1β not only in the various brain regions and plasma, as other studies have reported, but also in the CSF. The induction of IL1β in CSF is being reported for the first time in an animal model of psychological stress, which is consistent with findings in human patients of depression and anxiety [107,117–119].

With elevated neuroinflammation, we also replicate sexually divergent trends of behavioral impairment following physical and psychosocial stress in rats that we observed earlier with mouse model of stress. Female rats generally show a more pronounced phenotype for anxiety as well as neuroinflammation, which is also consistent with human data. For fear memory, we did not run a statistical comparison between males and females given the already complex ANOVA designs of our experiments, but once again females did seem to have a more robust phenotype as described in the results and the figure panels. In general, an elevation of IL1β is associated with impaired learning and memory *in vivo* [120–122], however, quite interestingly, we also observed that working memory deterioration was worse in male stressed rats. It is well understood that the neuroimmune response to any kind of injury, stress, or even neurodegeneration is not identical between the sexes in animal studies [123,124]. Moreover, working memory involves frontal areas of the brain that are distinct from hippocampus-amygdala circuits, which are mostly recognized for their role in fear and anxiety. It is quite plausible that there is some sex-specific plasticity that is selectively different in these brain areas that underlies the males showing more deficits in working memory vs. the females displaying more anxiety.

Additionally, this study contributes to the extant literature through our findings from pharmacological inhibition of canonical and non-canonical inflammasome mediators. Conceptually, the current literature indicates that both canonical and non-canonical inflammasome pathways can drive the IL1β response, thus stopping one pathway alone may not efficiently mitigate the IL1β response under stress. We therefore hypothesized that a combination of drugs blocking both canonical and non-canonical pathways that ultimately prevented both cleavage of pro-IL1β to IL1β and GSDMD to its N- and C-terminals would be a better strategy. Our investigation revealed that psychosocial stress triggers the activation of NLRP3-caspase 1 canonical inflammasomes as well as caspase 11-GSDMD non-canonical inflammasome mediators. Both mediators contribute to the surge in IL1β, which signals the development of behavioral dysfunction. To that end, we have provided robust evidence that inhibiting caspase 11 and GSDMD cleavage using wedelolactone and disulfiram, respectively, attenuates neuroinflammation and mitigates functional impairments. However, combination treatment of MCC950 (NLRP3/caspase 1 inhibitor; inhibiting the canonical inflammasome pathway) and wedelolactone (caspase 11 inhibitor; inhibiting the non-canonical inflammasome pathway) is more efficacious than individual treatment. Similarly, using psychologically stressed rats, we provided strong evidence that combination treatment with FDA-approved inhibitors ibrutinib (BTK-NLRP3-caspase 1/canonical inflammasome pathway inhibitor) and disulfiram (GSDMD cleavage/non canonical inflammasome pathway inhibitor) provided significantly better protection from neuroinflammation, dendritic spine elimination, and behavioral dysfunction. This is also a novel finding to our knowledge in an animal model of psychosocial stress. Our robust proof-of-concept preclinical preliminary evidence argues for considering this strategy for further investigation of potential pharmacotherapies for common psychological illnesses associated with stress, anxiety, and depression. To depict the involvement of canonical and non-canonical inflammasome mediators in psychosocial stress, we arrive at a model as illustrated in Figure 7J.

Our findings are also consistent with recent studies with disulfiram that have shown anxiolysis in mice [125]. Since disulfiram is routinely used by clinicians to treat alcohol abuse disorder because of its ability to inhibit alcohol dehydrogenase (ALDH), these authors also used a second ALDH inhibitor, cynamide, on their mouse model, but that did not have any anxiolytic effect. This also supports indirectly our interpretation that the reduction in anxious behavior we see from disulfiram treatment is a result of inhibition of GSDMD cleavage rather than its ALDH inhibiting properties.

We developed a cellular assay to validate the molecular mechanisms underlying peripheral inflammation following psychosocial stress. We isolated PBMCs from control and stressed rats and challenged them with vehicle or LPS to emulate stress. To decipher the underlying molecular pathway, we treated PBMCs with various canonical and non-canonical inhibitors. Our observation from studies with PBMCs was consistent with our studies from the brain samples from the rodent model of psychological stress. We demonstrated that inhibition of both canonical (MCC950 or ibrutinib) and non-canonical (wedelolactone or disulfiram) proved to be more efficacious in inhibiting IL1β release in culture media. To confirm the pyroptotic state of the immune cells, LPS-induced PBMCs were analyzed with SYTOX Green uptake, an indicator of pore formation in the plasma membrane. SYTOX Green uptake was significantly increased in LPS-challenged PBMCs from psychosocial stress. However, treatment with wedelolactone, ibrutinib, and disulfiram significantly reduced the uptake of SYTOX Green. In any of these groups, we did not observe significant differences with the WST-8, a cell viability assay, during the time frame of the experiment, indicating the absence of cell death. This suggests the development of a sub-lytic state, where the cell membrane is porous enough to allow the release of IL1β in culture media while maintaining the cell viability.

In conclusion, this study provides strong evidence that psychosocial stress induces both canonical and non-canonical inflammasome pathways, which jointly contribute to IL1β upregulation and a concomitant neuroinflammatory phenotype. The studies with the rodent social defeat stress model demonstrate that a combination treatment with both canonical and non-canonical inflammasome inhibitors proved more effective in mitigating behavioral deficits across multiple domains, including anxiety, social behavior, fear memory, and working memory. Novel findings include the non-canonical induction of caspase 11 and gasdermin D cleavage in brain and peripheral immune cells following psychosocial stress. While the present study illustrates the induction of non-canonical inflammasome mediators in psychosocial stress, it also raises many important questions, which can serve as the basis for further in-depth studies. We hope that our comprehensive set of experiments will form a stepping stone for further investigation on the involvement of different immune cells in the brain, such as microglia, astrocytes, macrophages, and infiltrating monocytes in the development of stress-induced neuroinflammation, blood-brain barrier impairment, synaptic pruning, and behavioral dysfunction. Further, a more in-depth study is required to decipher the molecular mechanisms underlying this stress-induced sub-lytic state of pyroptosis in immune cells. Overall, this study could also drive more granular investigations into the role of non-canonical inflammasomes in the development of peripheral inflammation in common psychiatric disorders.

## List of Abbreviations

ANOVA: Analysis of variance
ASC: Apoptosis-associated speck-like protein c
BSA: Bovine serum albumin
BTK: Bruton’s tyrosine kinase
CA3: Cornu ammonis 3
CCL2: C-C motif ligand 2
CCR2: C-C motif chemokine receptor 2
CSF: Cerebrospinal Fluid
CXCL2: C-X-C motif chemokine ligands
DMSO: Dimethyl sulfoxide
ELISA: Enzyme linked immunosorbent assay
EPM: Elevated plus maze test
GSDMD: Gasdermin D
i.g.: Intragastric gavage
i.p.: Intraperitoneal
ICAM1: Intercellular adhesion molecule-1
IL1β: Interleukin 1 Beta
LDT: Light–dark test
LPS: Lipopolysacharides
MCC950: 1,2,3,5,6,7-Hexahydro-s-indacen-4-ylcarbamoyl-[4-(2-hydroxypropan-2-yl)furan-2-yl]sulfonylazanide
MDD: Major depressive disorder
MGG: May-Grünwald Giemsa
NLRP3: Nucleotide-binding oligomerization domain (NOD) like receptor (NLR) family, pyrin domain containing protein 3
NOR: Novel object recognition
NS: Non-significant
OFT: Open field test
PAGE: Poly-acrylamide gel electrophoresis
PBMC: Peripheral blood mononuclear cells
PBS: Phosphate-buffered saline
PE-50: Polyethylene 50
PEG: Polyethylene glycol 400
p-NA: p-Nitroaniline
PTSD: Post-traumatic stress disorder
qRT-PCR: quantitative reverse transcription polymerase chain reaction
RFU: Relative fluorescent unit
RSD: Repeated social defeat
SD: Sprague Dawley
SDS: Sodium dodecyl sulfate
SSRI: Selective serotonin reuptake inhibitors
TBST: Tris-buffered saline with Tween

---

**Supplementary Figure 1.**
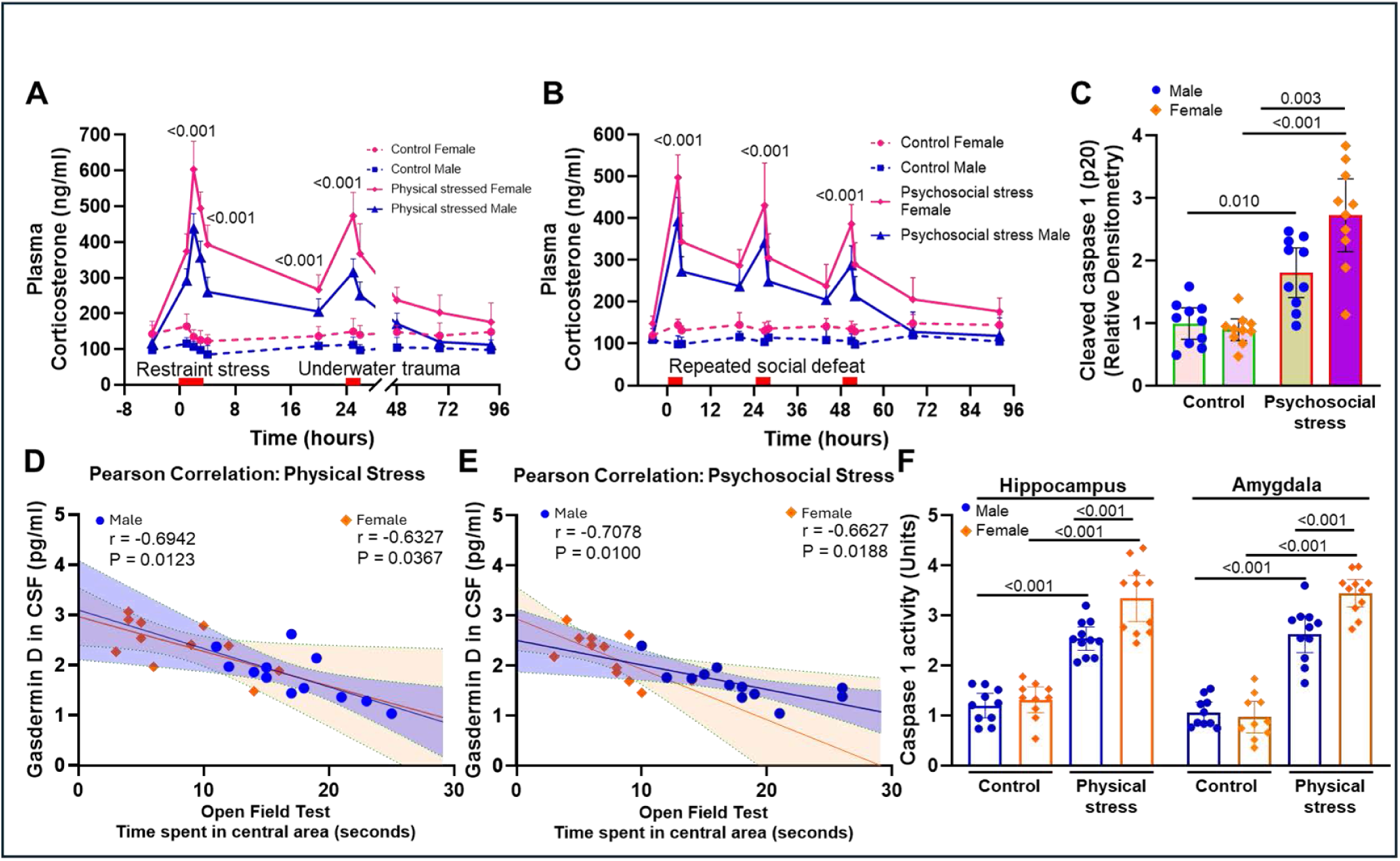
Physical and psychosocial stress induce corticosterone, caspase 1, and gasdermin. **D.** To evaluate the stress responses under physical **(A)** and psychosocial **(B)** stress in male and female rats, plasma corticosterone level was measured by the ELISA assay. Plasma corticosterone levels were significantly increased in females as compared to their male counterparts following physical and psychosocial stress. Immunoblot analysis from Figure 1 indicated induction of cleaved caspase 1 following psychosocial stress **(C).** Pearson correlation analysis was performed to determine the correlation of the gasdermin D levels in CSF with the time spent by rats in the central area of the open field test for rats with physical stress **(D)** and psychosocial stress **(E)**. Rats subjected to physical and psychosocial stress showed moderate negative correlation of the gasdermin D levels with the time spent in the open field test. Caspase 1 activity **(F)** was significantly higher in the hippocampus and amygdala of female stressed rats as compared to their male counterparts. All data are presented as mean with 95% CI (n=10-12); p-values are indicated for specific pairwise comparisons.

**Supplementary Figure 2.**
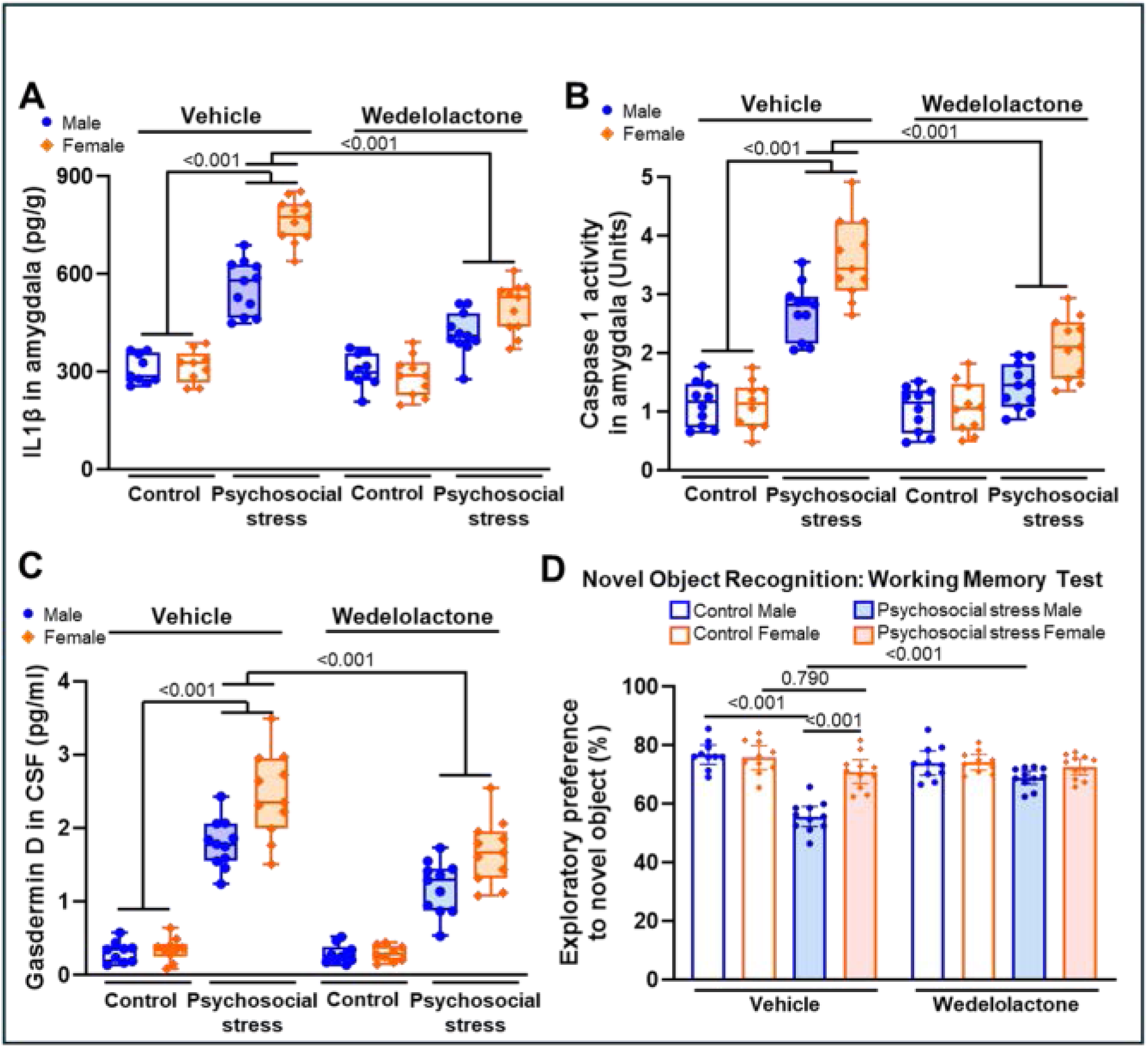
Wedelolactone treatment attenuates the induction of stress-induced IL1β, caspase 1, and gasdermin D. **(A)** Wedelolactone treatment significantly reduced IL1β levels in amygdala homogenates of stressed rats. Similarly, wedelolactone treatment also significantly reduced caspase 1 activity in the amygdala of stressed rats **(B)**. An ELISA assay indicated that the wedelolactone significantly reduced gasdermin D levels in CSF from rats with psychosocial stress **(C)**. Wedelolactone treatment significantly improved working memory, as indicated by increased exploration of the novel object in the novel object recognition test **(D)**. All data are presented as mean with 95% CI (n=10-12); p-values are indicated for specific pairwise comparisons.

**Supplementary Figure 3.**
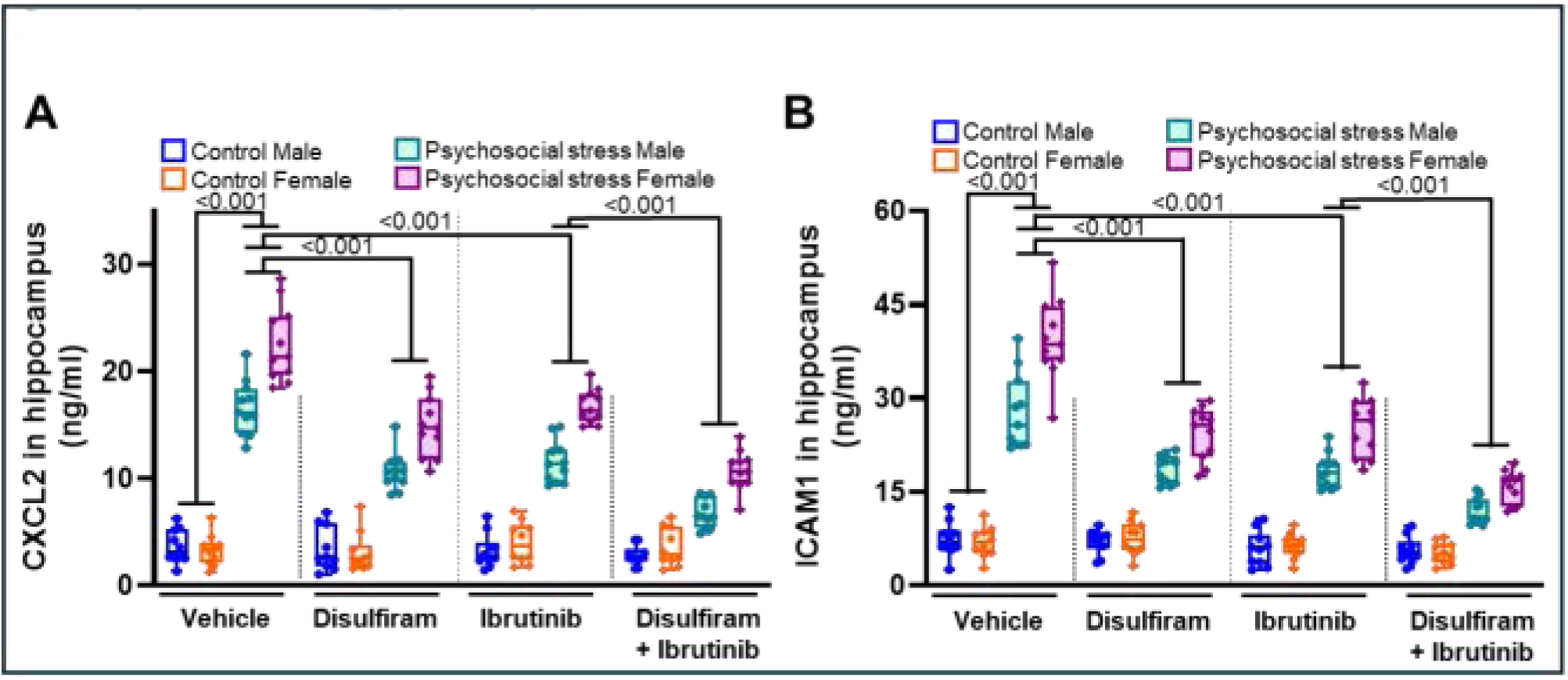
Psychosocial stress induces macrophage ligand CXCL2 and endothelial cell adhesion molecule ICAM-1 in the brain. The levels of macrophage ligand CXCL2 **(A)** and brain endothelial cell adhesion molecule ICAM-1 **(B)** were measured by ELISA in hippocampal homogenates. Female rats subjected to psychosocial stress showed significantly higher levels of CXCL2 and ICAM-1 in the hippocampus than their male counterparts. As compared to individual treatments, the combination treatment with disulfiram and ibrutinib significantly reduced the CXCL2 and ICAM-1 expressions. All data are presented as mean with 95% CI (n=10-12); p-values are indicated for specific pairwise comparisons.

**Supplementary Figure 4.**
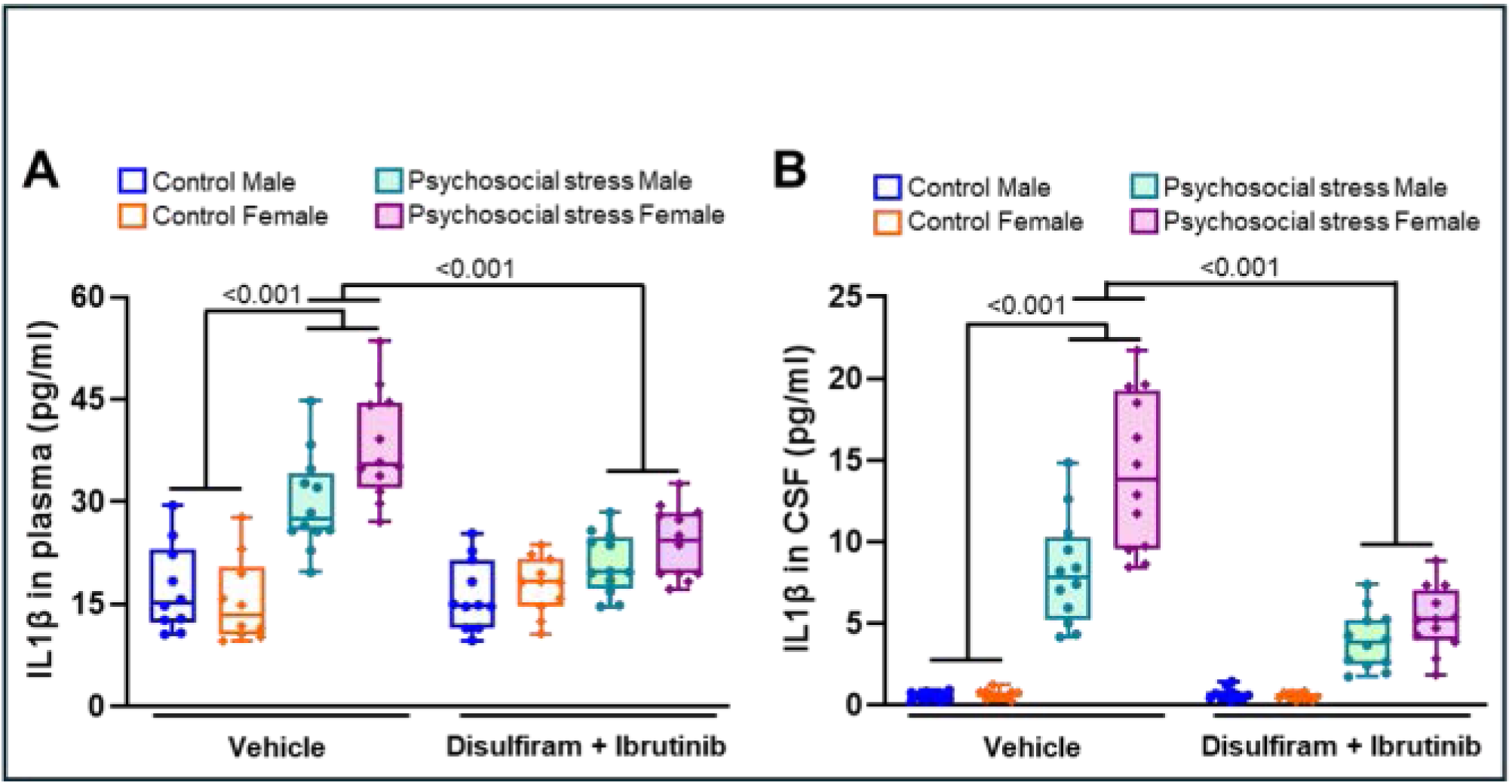
Combination treatment with disulfiram and ibrutinib attenuates IL1β levels in stressed rats. IL1β levels in plasma **(A)** and CSF **(B)** were measured by an ELISA assay. The combination treatment with disulfiram and ibrutinib significantly reduced IL1β levels in plasma and CSF of rats subjected to psychosocial stress. All data are presented as mean with 95% CI (n=10-12); p-values are indicated for specific pairwise comparisons.

**Supplementary Figure 5.**
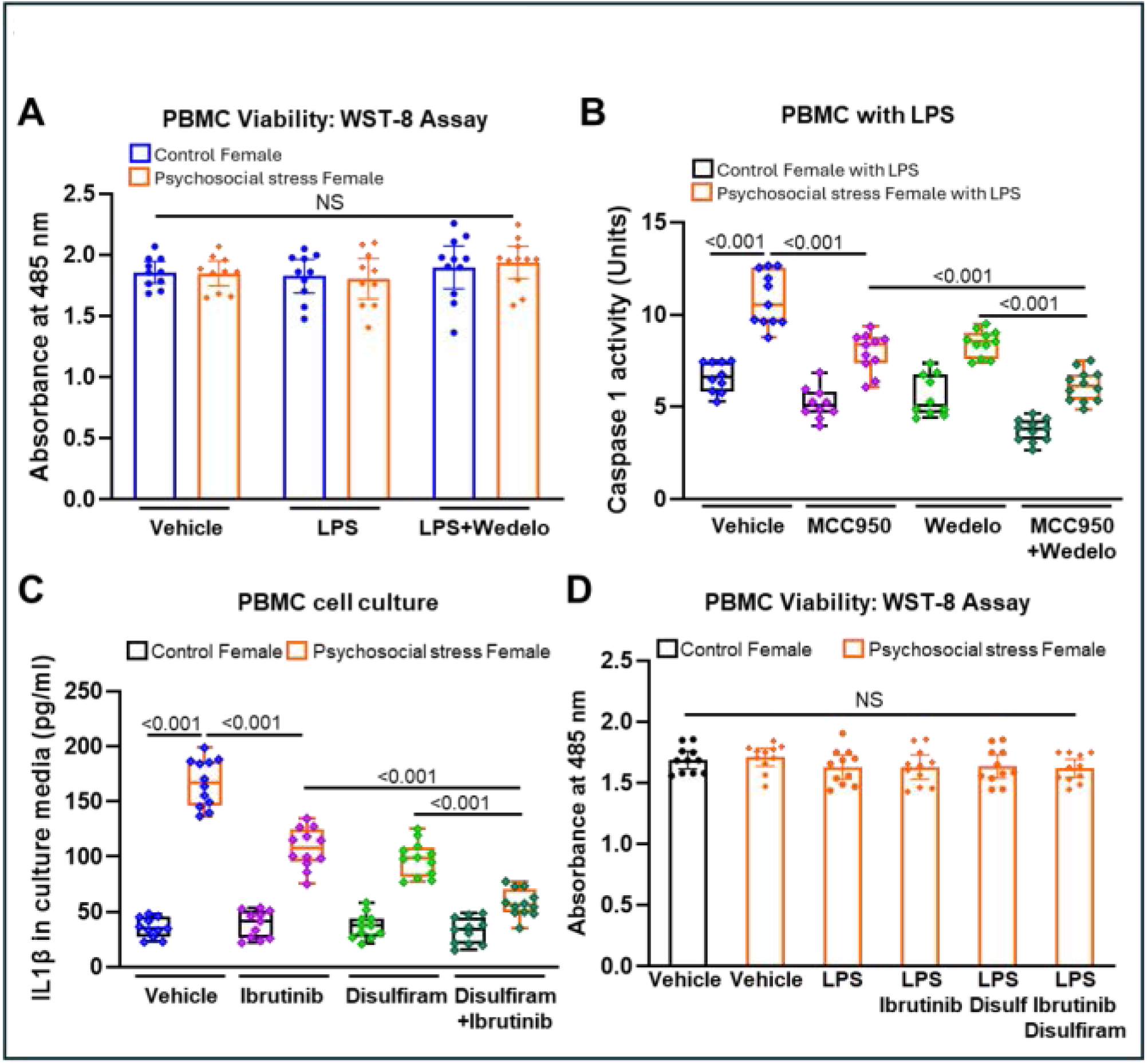
Inhibition of mediators of canonical and non-canonical inflammasomes suppresses IL1β and caspase 1 in PBMCs from stressed rats. PBMCs were isolated from control and stressed rats and then subjected to various treatments under *in vitro* conditions. IL1β levels in PBMC culture media were measured by an ELISA assay. As compared to individual treatments, a combination treatment with disulfiram and ibrutinib significantly and efficiently attenuated the IL1β levels in the culture media of PBMCs isolated from stressed rats (A). IL1β levels in PBMC culture media were significantly higher in psychosocial stress than in the control group. LPS-challenged PBMCs from psychosocially stressed female rats showed significantly higher caspase 1 activity (B). As compared to individual treatments, a combination treatment with MCC950 and wedelolactone significantly reduced the caspase 1 activity in LPS-challenged PBMCs from stressed rats. Cell viability was measured by WST-8 assay at 3 h following LPS challenge or treatment with ibrutinib and disulfiram (C-D). There were no significant differences in cell viability between various groups of PBMCs. All data are presented as mean with 95% CI (n=10-12); p-values are indicated for specific pairwise comparisons.

